# Acute manipulation and real-time visualization of membrane trafficking and exocytosis in *Drosophila*

**DOI:** 10.1101/2022.03.25.483021

**Authors:** Jade Glashauser, Carolina Camelo, Manuel Hollmann, Wilko Backer, Thea Jacobs, Jone Isasti Sanchez, Raphael Schleutker, Dominique Förster, Nicola Berns, Veit Riechmann, Stefan Luschnig

## Abstract

Intracellular trafficking of secretory proteins plays key roles in animal development and physiology, but tools for investigating dynamics of membrane trafficking have been limited to cultured cells. Here we present a system that enables acute manipulation and real-time visualization of membrane trafficking through reversible retention of proteins in the endoplasmic reticulum (ER) in living multicellular organisms. By adapting the “retention using selective hooks” (RUSH) approach to *Drosophila*, we show that trafficking of GPI-linked, secreted, and transmembrane proteins can be controlled with high temporal precision in intact animals and cultured organs. We demonstrate the potential of this approach by analyzing the kinetics of ER exit and apical secretion and the spatiotemporal dynamics of tricellular junction assembly in epithelia of living embryos. Furthermore, we show that controllable ER-retention enables tissue-specific depletion of secretory protein function. The system is broadly applicable to visualize and manipulate membrane trafficking in diverse cell types *in vivo*.

## Introduction

Secreted and membrane proteins comprise substantial portions of metazoan proteomes (Meinken et al., 2015; Pei et al., 2018). The secretory apparatus ensures the correct delivery of these proteins to the extracellular space, the cell surface or intracellular membrane compartments, where they carry out a wealth of functions in cell adhesion, cell shape regulation, motility, and signaling during development and homeostasis. Accordingly, mutations in many secretory pathway components cause developmental defects and diseases (Schotman and Rabouille, 2009; Yarwood et al., 2020). While many insights into the organization and function of the secretory pathway were obtained in cultured mammalian cells and in yeast, analogous studies in intact multicellular organisms have been limited by the lack of adequate tools to manipulate membrane trafficking *in vivo*. Analyzing the dynamics of these processes is challenging, because different pools of a given protein species move simultaneously along multiple intracellular routes (synthesis, exocytosis, endocytosis, recycling, degradation) and localize in different membrane compartments, which cannot be separated at steady state. Therefore, the synthesis or transport of selected proteins needs to be synchronized to follow their routes through the secretory apparatus. Classical methods for synchronizing secretory protein trafficking either rely on temperature blocks (Griffiths et al., 1985) or drugs (e.g., Brefeldin A; Lippincott-Schwartz et al., 1989) to reversibly arrest intracellular transport, or employ special conditionally mis-folded or aggregated proteins that are retained in the endoplasmic reticulum (ER). Release of these proteins from the ER is achieved by shifting cells to permissive temperature (Kreis and Lodish, 1986; Lafay, 1974), by adding a small-molecule ligand (Casler et al., 2020; Rollins et al., 2000), or by illumination with UV light (Chen et al., 2013). Although these approaches have revealed fundamental insights into the organization and dynamics of the secretory apparatus, they are limited to special proteins (*e*.*g*., the conditional thermosensitive mutant viral glycoprotein VSVGtsO45; Kreis and Lodish, 1986; Presley et al., 1997; Scales et al., 1997) and require treatments using non-physiological temperatures, drugs, or potentially damaging doses of UV light.

A powerful experimental system that avoids these limitations is the ‘*retention using selective hooks’* (RUSH) system (Boncompain et al., 2012). RUSH enables synchronization of trafficking by using a two component-system comprising (i) a secretory cargo protein fused with a fluorophore and a streptavidin-binding peptide (SBP) tag and (ii) a streptavidin (SA) “hook” protein targeted to a membrane “donor” compartment of choice by a signal sequence (*e*.*g*., a KDEL motif for retention in the ER). SBP-Cargo and SA-hook proteins form a complex that is retained in the donor compartment (Fig. 1A). Addition of biotin rapidly dissociates the cargo-hook complex, triggering release of cargo from the donor compartment, and allows to follow the synchronized passage of the fluorescent cargo molecules through the secretory apparatus. Importantly, RUSH is applicable to diverse cargo proteins, and their trafficking can be controlled at physiological temperatures using a non-toxic cell-permeable small molecule. This approach revealed important insights into the molecular events underlying ER exit (Shomron et al., 2021), ER-to-Golgi transport (Weigel et al., 2021; Westrate et al., 2020), intra-Golgi trafficking, and transport of cargo between the Golgi apparatus and the plasma membrane (Fourriere et al., 2016; Stalder and Gershlick, 2020). However, applications of the RUSH system have thus far been limited to cultured cells, which typically do not represent the complexity of tissues in multicellular organisms, where membrane trafficking is developmentally or physiologically regulated.

**Figure 1.**
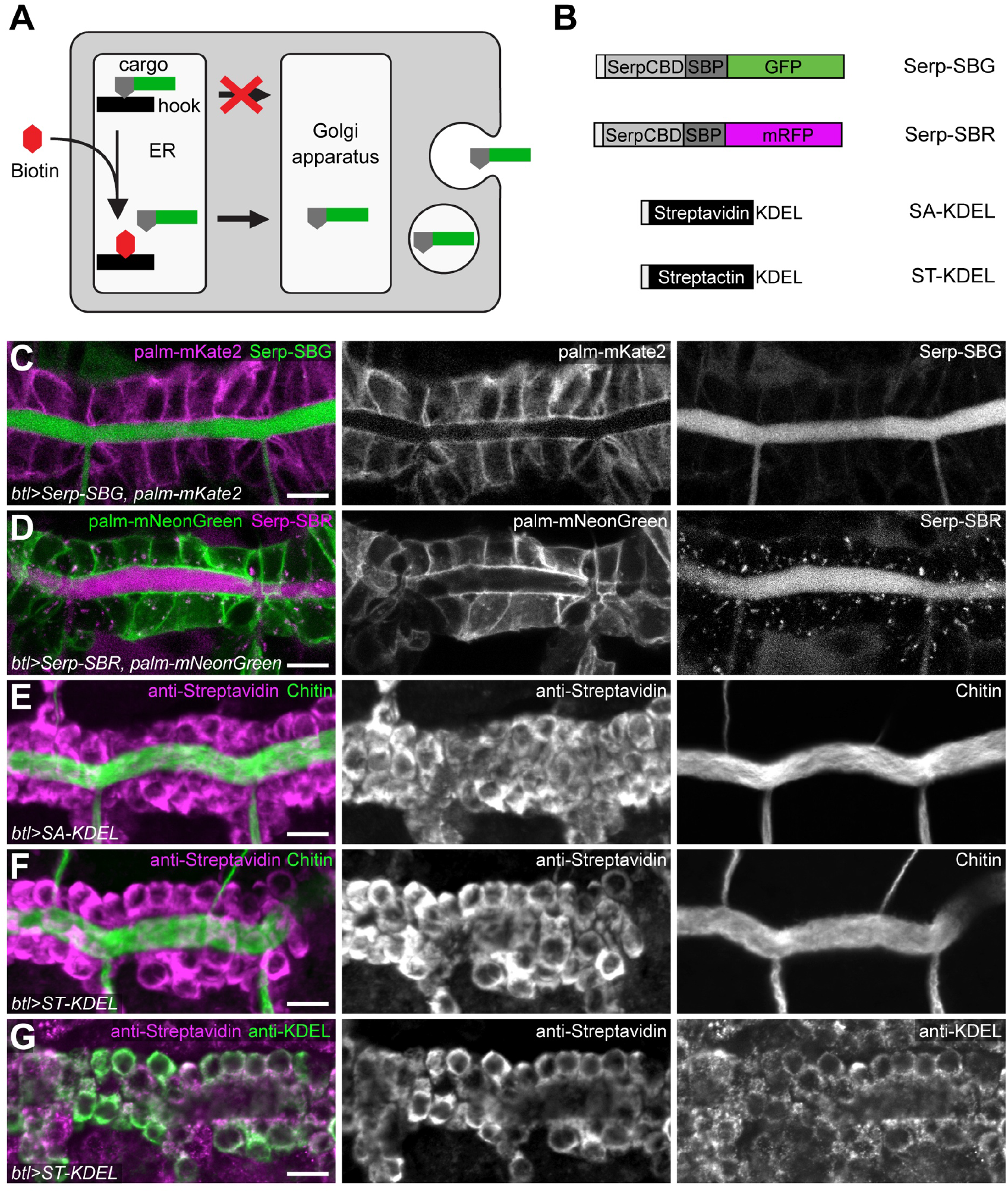
A two-component system for controlling secretory protein trafficking *in vivo*. **(A)** Principle of RUSH. A streptavidin “hook” protein carrying an ER-retention signal is retained in the ER lumen and binds a secretory cargo protein tagged with a fluorescent protein and a Streptavidin-binding peptide (SBP), leading to ER retention of the cargo. Addition of biotin dissociates the hook-cargo complex, thus allowing cargo to exit the ER and synchronizing the passage of cargo through the secretory apparatus. **(B)** Secretory cargo and ER-resident “hook” constructs. Cargo proteins consist of the N-terminal signal peptide and chitin-binding domain of Serpentine (Serp-CBD) fused to a Streptavidin-binding peptide (SBP) and a C-terminal EGFP (Serp-SBG, green) or mRFP (Serp-SBR, magenta) tag. ER-resident hook proteins comprise either core streptavidin (SA) or Streptactin (ST) fused to an N-terminal signal peptide and a C-terminal KDEL sequence. **(C**,**D)** Confocal sections of tracheal dorsal trunk (metamere seven) in living embryos expressing Serp-SBG (green; C) and palmitoylated mKate2 (palm-mKate2; magenta; C) or Serp-SBR (magenta; D) and palmitoylated mNeonGreen (palm-mNeonGreen; green, D) in tracheal cells under control of *btl-*Gal4. Anterior is to the left and dorsal is up here and in all subsequent panels, unless noted otherwise. **(E**,**F)** Confocal sections of tracheal dorsal trunk in fixed embryos expressing streptavidin-KDEL (SA-KDEL; magenta, E) or Streptactin-KDEL (ST-KDEL; magenta, F) in tracheal cells, stained with antibodies against streptavidin (magenta) and for chitin (tracheal lumen, green). **(G)** Confocal section of tracheal dorsal trunk in embryo expressing Streptactin-KDEL, stained with anti-streptavidin (magenta) and anti-KDEL (green) antibodies to label ER. Note localization of ST-KDEL in the ER. Scale bar: (C-G), 5 μm.

We present a set of tools that enable synchronization and visualization of membrane trafficking in intact animals and in cultured organs through reversible ER-retention of secretory proteins. The system is broadly applicable to investigate the spatiotemporal dynamics of secretory trafficking during development *in vivo*. Additionally, hook-induced ER retention provides a powerful approach to deplete membrane protein function in a tissue-specific manner.

## Design

### A two-component system for synchronization of membrane trafficking *in vivo*

To control trafficking of secretory proteins in *Drosophila*, we generated a set of cargo and hook proteins (Fig. 1B, Table S1) that can be expressed in a tissue-specific manner using the Gal4/UAS system (Brand and Perrimon, 1993). As a model cargo protein we chose the chitin deacetylase Serpentine (Serp), which is secreted by embryonic tracheal cells into the tube lumen (Luschnig et al., 2006; Wang et al., 2006). The N-terminal portion of Serp comprising the signal peptide and chitin-binding domain (CBD) is sufficient to direct apical secretion of a Serp(CBD)-GFP fusion protein, resembling the trafficking of full-length Serp-GFP protein, but unlike full-length Serp-GFP, expression of Serp(CBD)-GFP does not cause tracheal defects (Luschnig et al., 2006; Wang et al., 2006). Therefore, to generate model cargo proteins for RUSH experiments, we fused the N-terminal portion of Serp to Streptavidin-binding peptide (SBP, 38 aa; Keefe et al., 2001) followed by either the EGFP or mRFP coding sequence, resulting in Serp(CBD)-<u>SB</u>P-<u>G</u>FP (Serp-SBG) and Serp(CBD)-<u>SB</u>P-m<u>R</u>FP (Serp-SBR), respectively (Fig. 1B). When expressed in tracheal cells under the control of *btl*-Gal4, Serp-SBG and Serp-SBR accumulated in the tracheal lumen (Fig. 1C,D), resembling endogenous Serp protein (Luschnig et al., 2006; Wang et al., 2006). To retain the cargo proteins in the ER, we generated ER-resident hook constructs comprising either core streptavidin (SA) or the SA variant Streptactin (ST; Voss and Skerra, 1997) fused to the N-terminal signal peptide of Serp and a C-terminal ER retention signal (KDEL; Fig. 1B). When expressed in tracheal cells, SA-KDEL and ST-KDEL were distributed in a perinuclear pattern (Fig. 1E,F) and co-localized with anti-KDEL immunostaining (Fig. 1G), indicating localization in the ER.

## Results

### SBP-tagged proteins can be retained in the ER by Streptavidin hooks

To test whether streptavidin hooks can retain Serp-SBG in the ER, we co-expressed SA-KDEL and Serp-SBG in embryonic tracheal cells. However, expression of a single-copy UAS-SA-KDEL transgene did not modify the distribution of Serp-SBG, which was secreted into the tracheal lumen like in control embryos not expressing SA-KDEL (Fig. 2A,B). Since SA forms a tetramer with two SBP binding sites (Barrette-Ng et al., 2013), we reasoned that an excess of SA-KDEL is required for efficient retention of Serp-SBG. Therefore, to increase SA-KDEL levels, we used a vector (pUASTΔSV40; Nelson et al., 2018) that yields approximately 5-fold higher expression levels compared to the pUAST vector (Brand and Perrimon, 1993) used for the first generation of SA-KDEL constructs. Indeed, combining one copy of UASTΔSV40-SA-KDEL (referred to as UAS-SA-KDEL(strong)) with one copy of UAS-Serp-SBG led to partial intracellular accumulation of Serp-SBG (Fig. 2C). To improve retention efficiency, we combined multiple copies of UAS-SA-KDEL(strong) transgenes with a single copy of UAS-Serp-SBG (Fig. 2C-E). The ratio of intracellular to luminal Serp-SBG signals increased with the dosage of SA-KDEL hooks, and maximal retention (median ratio of 1.2; n=15) was observed in 95% of embryos (n=40) expressing four copies of SA-KDEL(strong) (Fig. 2E,F). We generated strains carrying two insertions of UAS-SA-KDEL(strong) or UAS-ST-KDEL(strong), respectively, for each chromosome, enabling straightforward and versatile use in genetic crosses (Table S1; STAR Methods).

**Figure 2.**
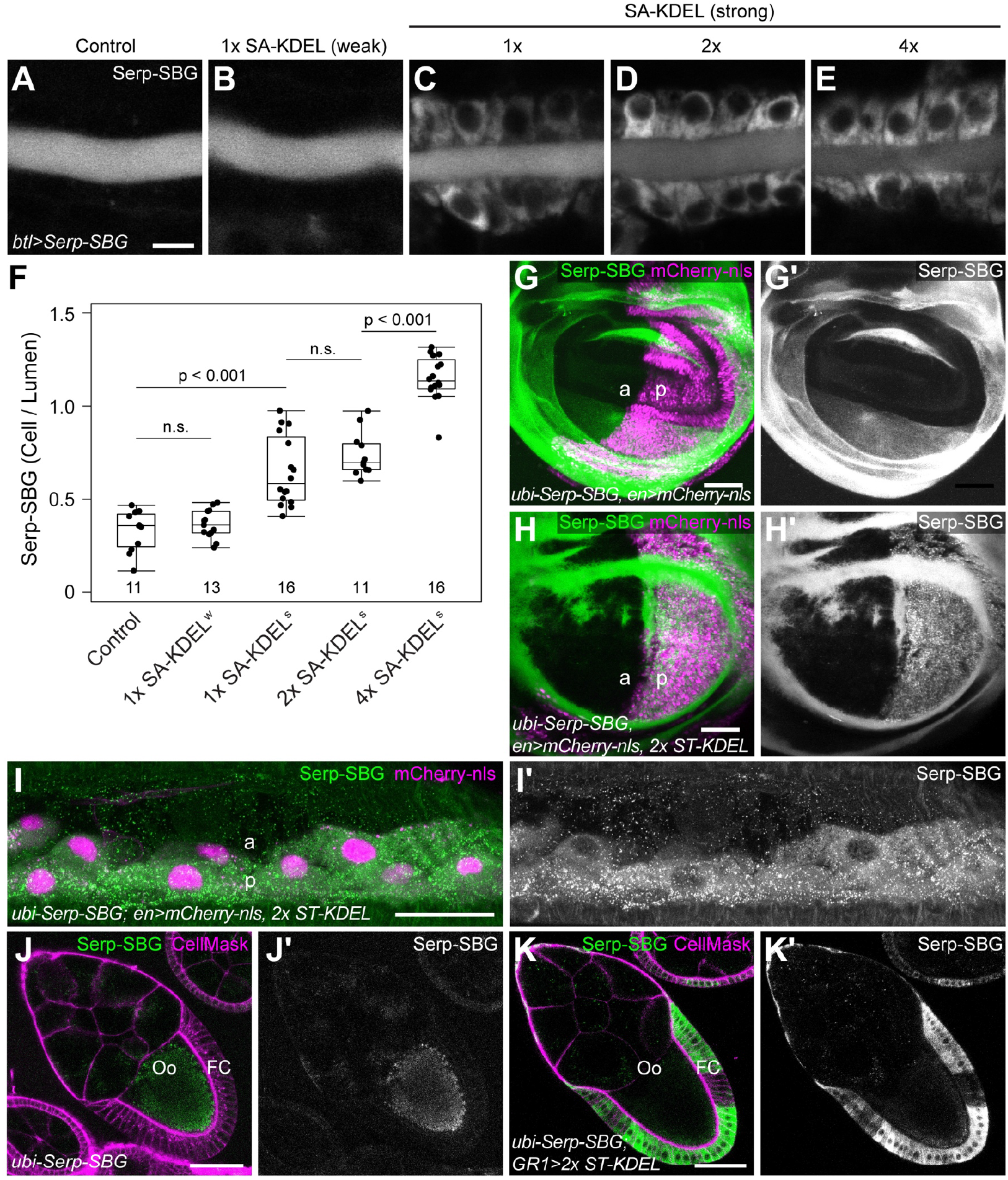
The efficiency of ER retention depends on the ratio of cargo to hook proteins. **(A-E)** Confocal sections of dorsal trunk in living embryos (stage 15) expressing Serp-SBG alone (control; A), with one copy of SA-KDEL(weak) (B), or with the indicated number of copies of SA-KDEL(strong) (C-E) in tracheal cells. Note that intracellular Serp-SBG signals increase, and luminal signals decrease with increasing dosage of SA-KDEL. **(F)** Quantification of the ratio of intracellular and luminal Serp-SBG signals. Number of embryos analyzed per genotype is indicated. Boxplot shows median (line), interquartile range (box) and 1.5x interquartile range from the 25th and 75th percentile (whiskers). p-values (Student’s t-test) are indicated; n.s., not significant. **(G-H’)** Wing imaginal discs of third-instar larvae expressing Serp-SBG (green) ubiquitously under control of the *ubiquitin* (*ubi*) promoter. mCherry-nls (magenta) expression under control of *en*-Gal4 marks the posterior compartment. In control disc (G,G’), Serp-SBG accumulates in the disc lumen. Expression of ST-KDEL (two copies) in the posterior compartment leads to intracellular retention of Serp-SBG in ST-KDEL-expressing cells (H,H’). a, anterior compartment; p, posterior compartment. **(I**,**I’)** Epidermis of third-instar larvae expressing Serp-SBG (green) ubiquitously as in (G,H). *en*-Gal4 drives expression of mCherry-nls (magenta) and ST-KDEL (two copies) in posterior (p) compartment cells. Note diffuse distribution of Serp-SBG in cuticle of anterior (a) compartment and intracellular retention in posterior compartment cells. Anterior is up. **(J-K’)** Ovarian egg chambers (stage 10A) from adult females expressing Serp-SBG (green) ubiquitously under control of the *ubi* promoter. CellMask (magenta) marks plasma membranes. In control egg chamber (J,J’), Serp-SBG is secreted by follicle cells (FC) and taken up into the oocyte (Oo). Expression of ST-KDEL (two copies) in follicle cells driven by GR1-Gal4 leads to intracellular retention of Serp-SBG (K,K’). Anterior is to upper left. Scale bars: (A-E), 5 μm; (G-K), 50 μm.

To ask whether hook-mediated ER retention works in different tissues and developmental stages, we generated flies expressing Serp-SBG ubiquitously under the control of the *ubiquitin* promoter (*ubi*-Serp-SBG) and used tissue-specific Gal4 drivers to express SA-KDEL or ST-KDEL hooks in selected tissues. In wing imaginal discs of third-instar larvae, Serp-SBG was secreted into the disc lumen (Fig. 2G). Expression of SA-KDEL or ST-KDEL hooks in the posterior compartment using *engrailed-*Gal4 (*en*-Gal4) led to intracellular retention of Serp-SBG in the hook-expressing cells (Fig. 2H). Likewise, Serp-SBG was retained in hook-expressing posterior compartment cells in the larval epidermis (Fig. 2I). Intracellular retention of Serp-SBG was also observed in adult tissues, such as ovarian follicle cells expressing the ER hooks (Fig. 2J,K). Thus, SA-KDEL and ST-KDEL hooks mediate efficient ER retention of an SBP-tagged secreted protein in different tissues throughout development.

We wondered whether elevated secretory cargo load caused by long-lasting expression of streptavidin hooks or by hook-mediated ER-retention of secreted proteins might induce ER stress and consequently activate the unfolded protein response (UPR; Hetz et al., 2020; Ryoo, 2015). To test for UPR induction, we used an Xbp1-GFP reporter construct, which yields a nuclear-localized Xbp1-GFP fusion protein only after unconventional splicing of Xbp1-GFP mRNA mediated by the nuclease IRE-1 in response to ER stress (Ryoo et al., 2007). Driving Xbp1-GFP expression in tracheal cells of control embryos did not result in nuclear Xbp1-GFP signals (Fig. S1A,K), whereas nuclear Xbp1-GFP was induced upon RNAi-mediated knockdown of the ER chaperone Heat shock protein 70 cognate 3 (Hsc70-3; Fig. S1E,K), which is required for protein folding and quality control in the ER (Chow et al., 2015). Nuclear Xbp1-GFP signals were not detectable in tracheal cells expressing two copies of SA-KDEL (Fig. S1B,K), but appeared in a subset of cells expressing four copies of SA-KDEL (Fig. S1C,K), suggesting that SA-KDEL can induce ER stress in a dosage-dependent manner. However, this was not associated with morphological abnormalities or cell death. Xbp1-GFP levels did not increase when Serp-SBR was retained by SA-KDEL in the ER of tracheal cells (Fig. S1D,K), suggesting that ER retention of SBP-tagged cargo does not induce ER stress. In ovarian follicle cells, nuclear Xbp1-GFP was not detectable in the presence of either two or four copies of SA-KDEL, and only rare nuclei showed Xbp1-GFP signals when Serp-SBR was retained in the ER (Fig. S1F-I, L), whereas treatment with the reducing agent Dithiothreitol (DTT) led to Xbp1-GFP accumulation throughout the follicle epithelium (Fig. S1J,L). Thus, while high levels of SA-KDEL expression can induce ER stress in tracheal cells, ER stress was not induced in follicle cells, consistent with the notion that the sensitivity to ER cargo load differs between cell types (Sone et al., 2013).

### Injection of biotin triggers ER exit and secretion of SBP-tagged cargo

To test whether SA-KDEL-mediated ER retention is reversible, we injected biotin (1 mM) into the hemocoel of embryos (stage 15) expressing Serp-SBG (one copy) and SA-KDEL (four copies) in tracheal cells (Fig. 3). We analyzed the distribution of Serp-SBG in the multicellular dorsal trunk (DT; Fig. 3A,B, Movie S1, Movie S2) and in unicellular dorsal branches (Fig. 3C, Movie S3). Before biotin injection, Serp-SBG was distributed in a perinuclear pattern in tracheal cells. Approximately two minutes after biotin injection, Serp-SBG started to accumulate in discrete intracellular puncta. Six to ten minutes after injection, Serp-SBG signals in the tracheal lumen started to increase rapidly, while intracellular signals concomitantly decreased (Fig. 3A,C,D, Movie S1), indicating that Serp-SBG passed the secretory pathway and started to be secreted as early as six minutes after release from the ER. Luminal Serp-SBG signals were maximal after 30 minutes and then declined gradually, while intracellular signals concurrently leveled off (Fig. 3D), suggesting that the supply of ER-retained Serp-SBG protein was depleted. The kinetics of ER release and luminal secretion of Serp-SBG were dependent on the concentration of injected biotin. 1 mM biotin induced rapid ER release, whereas 10 μM biotin was not sufficient to induce release (n=3; Fig. S2, Movie S4). A slow and steady increase in luminal Serp-SBG signals upon injection of 10 μM biotin, but no concomitant reduction of intracellular signals, suggests that constant slow leakage of Serp-SBG from the ER is compensated by newly synthesized Serp-SBG protein (Fig. S2).

**Figure 3.**
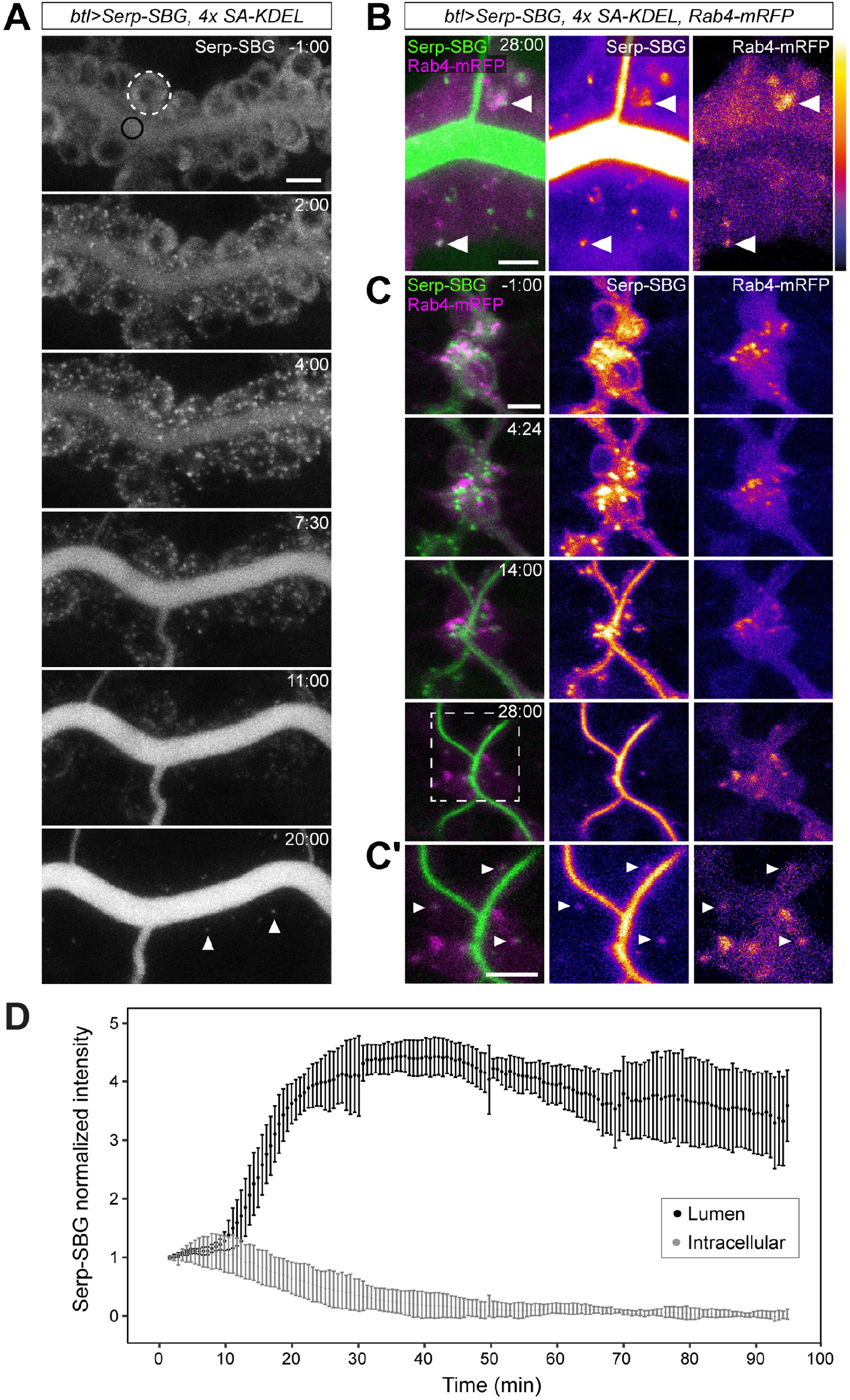
Biotin injection triggers rapid ER exit and exocytosis of Serp-SBG in tracheal cells. **(A)** Stills (maximum-intensity projections) from time-lapse movie of embryo (stage 15) expressing Serp-SBG and SA-KDEL (four copies) in tracheal cells. After biotin injection (1 mM) at t=0 min, Serp-SBG is released from ER, begins to accumulate at immobile puncta (ER exit sites) 2 min after injection, and is subsequently secreted into the lumen. Rapidly moving intracellular Serp-SBG-positive puncta, corresponding to endosomes, appear 20 min after injection (arrowheads). ROIs in tracheal cells (dashed white circle) and lumen (black circle) are indicated in top panel. Time (min:s) is indicated. See Movie S1. **(B)** Confocal section of dorsal trunk in embryo (stage 15) expressing Serp-SBG, Rab4-mRFP and SA-KDEL (four copies) in tracheal cells, 28 min after biotin injection. A subset of intracellular Serp-SBG puncta overlaps with recycling endosomes marked by Rab4-mRFP (arrowheads). Single-channel images display Serp-SBG and Rab4-mRFP intensities as a heat map. Time (min:s) is indicated. See Movie S2. **(C**,**C’)** Stills (maximum-intensity projections) of dorsal branch from time-lapse movie of embryo (stage 16) expressing Serp-SBG, Rab4-mRFP and SA-KDEL (four copies) in tracheal cells. After injection of biotin at t=0 min, Serp-SBG is rapidly secreted into the lumen. At t=28 min, Serp-SBG is detectable in recycling endosomes marked by Rab4-mRFP (C’, arrowheads). Anterior is to the left. Time (min:s) is indicated. See Movie S3. **(D)** Quantification of Serp-SBG intensity in the lumen (black) and inside tracheal cells (grey) of dorsal trunk after biotin injection at t=0 min. Mean intensities ± s.d., normalized to intensities at t=0 min, are shown. n=5 embryos. Scale bars: (A,B,C,C’), 5 μm.

From approximately 20 minutes after injection, Serp-SBG appeared in small intracellular puncta (0.67 ± 0.12 μm diameter, mean ± s.d., n=15 puncta), which moved rapidly (0.43 ± 0.18 μm/s; mean ± s.d., n=22 puncta) and co-localized partially with Rab4-mRFP-positive recycling endosomes (Fig. 3B,C’, Movie S2, Movie S3). This suggests that secreted Serp-SBG was internalized from the lumen and entered the endocytic pathway, consistent with intense endocytic activity of tracheal cells during luminal clearance (Dong et al., 2013; Tsarouhas et al., 2007). Taken together, these findings show that biotin injection triggers rapid release of retained proteins from the ER, enabling synchronization and visualization of secretory protein trafficking in living embryos.

### Serp-SBG rapidly passes the secretory apparatus

We used this system to analyze the dynamics of Serp-SBG transport through the early secretory apparatus. Embryonic tracheal cells contain on the order of a dozen dispersed Golgi stacks (Armbruster and Luschnig, 2012; Förster et al., 2010), each of which resides adjacent to a transitional ER (tER; Schweizer et al., 1990) unit, where secretory cargo proteins are collected at clusters of ER exit sites (ERES; Bannykh et al., 1996) before leaving the ER. We labeled ERES using an mCherry-tagged version of the COPII component Sec24AB (mCherry-Sec24; Fig. 4, Fig. S3), which colocalized with the ERES marker Sec16 (Fig. S3A; Ivan et al., 2008) and localized adjacent to the *cis*-Golgi marker GRASP65-GFP (Fig. S3B; Yang et al., 2021).

**Figure 4.**
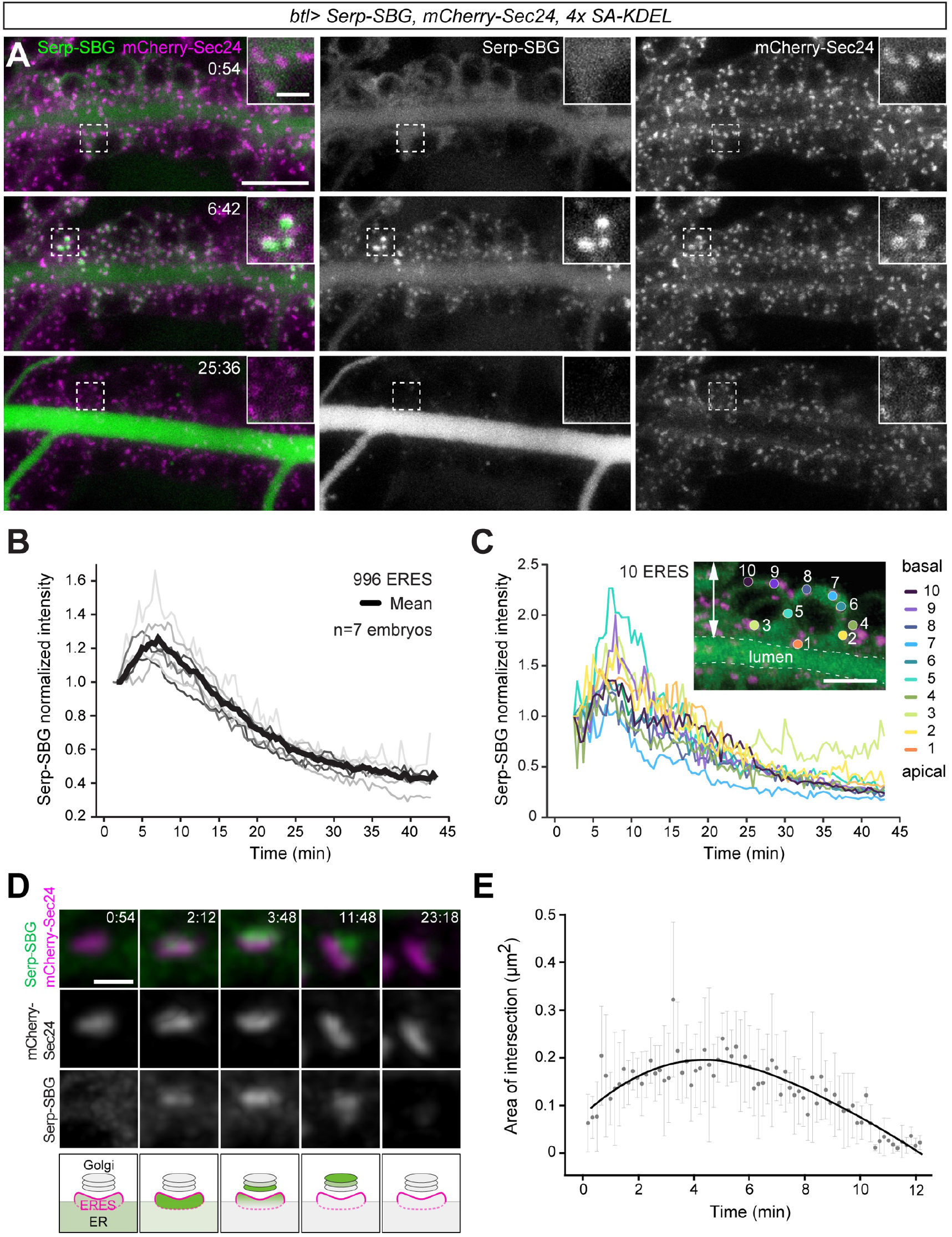
Rapid ER exit and passage of the Golgi apparatus by Serp-SBG. **(A)** Stills from time-lapse movie of tracheal dorsal trunk in embryo (stage 15) expressing Serp-SBG, mCherry-Sec24 and four copies of SA-KDEL in tracheal cells. Biotin was injected at t=0 min. Serp-SBG (green) accumulates at ER exit-sites (mCherry-Sec24, magenta) at t=6:42. Time (min:s) is indicated. See Movie S5. **(B)** Quantification of Serp-SBG signals at mCherry-Sec24-positive ERES. Each thin line indicates the mean of Serp-SBG intensity of at least 26 ERES per time point in one embryo, thick black line shows the mean of seven embryos. Data were normalized to mean intensities at the first time point. Serp-SBG accumulation at ERES is maximal seven minutes after biotin injection. **(C)** Quantification of Serp-SBG levels at ten ERES distributed from apical (1) to basal (10) in tracheal cells indicates similar profiles of Serp-SBG trafficking. **(D)** High-resolution images showing Serp-SBG (green) at a single mCherry-Sec24-labeled ERES (magenta) after biotin injection at t=0 min. After ER release, Serp-SBG colocalizes with mCherry-Sec24 (t=2:12) and subsequently shifts next to the mCherry-Sec24-marked ERES, presumably residing in the Golgi apparatus (t=3:48, 11:48), before disappearing from the ERES-Golgi complex (t=23:18). Images were processed using deconvolution. See Movie S6. Scheme (bottom panel) illustrates passage of Serp-SBG (green) through the ERES (magenta) and Golgi apparatus (grey). See Movie S6. **(E)** Quantification of the area of intersection between Serp-SBG and mCherry-Sec24. Each data point represents the mean ± s.d. of the overlapping area between Serp-SBG and mCherry-Sec24 signals. Between 38 and 58 mCherry-Sec24-positive ERES were analyzed per time point. The black curve shows a local polynomial regression fit. Scale bars: (A), 10 μm, insets: 2 μm; (C), 5 μm; (D) 1 μm.

To analyze the transit of Serp-SBG through the ER-Golgi interface, we measured Serp-SBG signals within spherical volumes, each of which enclosed an mCherry-Sec24-labelled ERES (Fig. 4A,B; Movie S5). Shortly after biotin injection, Serp-SBG rapidly accumulated at mCherry-Sec24-labelled ERES in a synchronous fashion with a peak at seven minutes after injection, and subsequently decreased over 30 minutes (Fig. 4B; n=996 mCherry-Sec24 units in 7 embryos). Afterwards, Serp-SBG levels at ERES leveled off, suggesting that most of the retained Serp-SBG protein has exited the ER. Comparable profiles of Serp-SBG levels were detected in all mCherry-Sec24-labelled tER units analyzed, independent of their position along the apical-basal axis of tracheal cells (Fig. 4C), suggesting that Serp-SBG passes through all ERES/Golgi units in a uniform fashion.

High-resolution imaging of individual ERES/Golgi units revealed that Serp-SBG started to accumulate at ERES, as detected by colocalization with mCherry-Sec24, approximately two minutes after biotin injection (Fig. 4D,E; Movie S6). Shortly after, the bulk of Serp-SBG shifted into clusters adjacent to and partially overlapping with the mCherry-Sec24-labelled ERES, before disappearing approximately 20 minutes after biotin injection (Fig. 4D, Movie S6). The area of overlap between mCherry-Sec24 and Serp-SBG signals was maximal between two and six minutes and then dropped to zero approximately 12 minutes after biotin injection (Fig. 4E). These findings suggest that after residing in the tER or at ERES for approximately four minutes, the bulk of Serp-SBG translocates to an adjacent compartment, corresponding to Golgi cisternae (Yang et al., 2021), where the protein spends approximately eight minutes before exiting the Golgi apparatus (Fig. 4E). Thus, RUSH allows analyzing the dynamics of trafficking between different membrane compartments *in vivo*.

### RUSH enables control over trafficking of endogenous proteins

Having shown that secretion of Serp-SBG can be controlled using SA hooks, we asked whether the system is also applicable to endogenous proteins. To this aim, we took advantage of the Cambridge protein trap insertion (CPTI; Lowe et al., 2014; Lye et al., 2014) fly strains, in which a given protein is tagged with Venus-YFP and two flanking StrepII tags (<u>S</u>trepII-<u>V</u>enus-YFP-<u>S</u>trepII; SVS) encoded by an artificial exon inserted into the endogenous gene locus (Fig. 5, Fig. S4). We tested transmembrane (Basigin (Bsg), Echinoid (Ed), Gliotactin (Gli), Neurexin 4 (Nrx4), Notch (N), Sidekick (Sdk)), Glycosylphosphatidylinositol (GPI)-anchored (Fasciclin 2 (Fas2), Lachesin (Lac)), and secreted (Chitin deacetylase-like 4; Cda4) proteins carrying an SVS tag in their extracellular portion (Fig. S4). To test whether localization of these proteins can be manipulated using ER hooks, we drove expression of ER hooks in epidermal stripes (using *en*-Gal4 or *hh*-Gal4) in heterozygous animals carrying one copy of the SVS-tagged locus (Fig. 5, Fig. S4). To maximize the affinity for StrepII-tagged proteins, we used Streptactin (ST)-KDEL instead of SA-KDEL hook proteins, since the StrepII tag binds with higher affinity to Streptactin (ST-StrepII: K_d_ = 10^−6^ M; Voss and Skerra, 1997) than to streptavidin (SA-StrepII: K_d_ = 72 × 10^−6^ M; Kim et al., 2010). Expression of two copies of ST-KDEL indeed led to ER retention of SVS-tagged proteins, as shown for Fas2::SVS in the embryonic and larval epidermis, larval imaginal discs, and in adult ovarian follicle cells (Fig. 5A-F). While efficient ER retention was observed for the majority of SVS-tagged proteins tested (Cda4, Fas2, Gli, Notch, Nrx4; Fig. S4), some proteins showed partial (Lac; Fig. S4H) or no detectable retention (Bsg, Ed, Sdk; Fig. S4D-F), suggesting that cargo protein abundance or accessibility of the SVS tag can influence the efficiency of ST-KDEL-induced ER retention.

**Figure 5.**
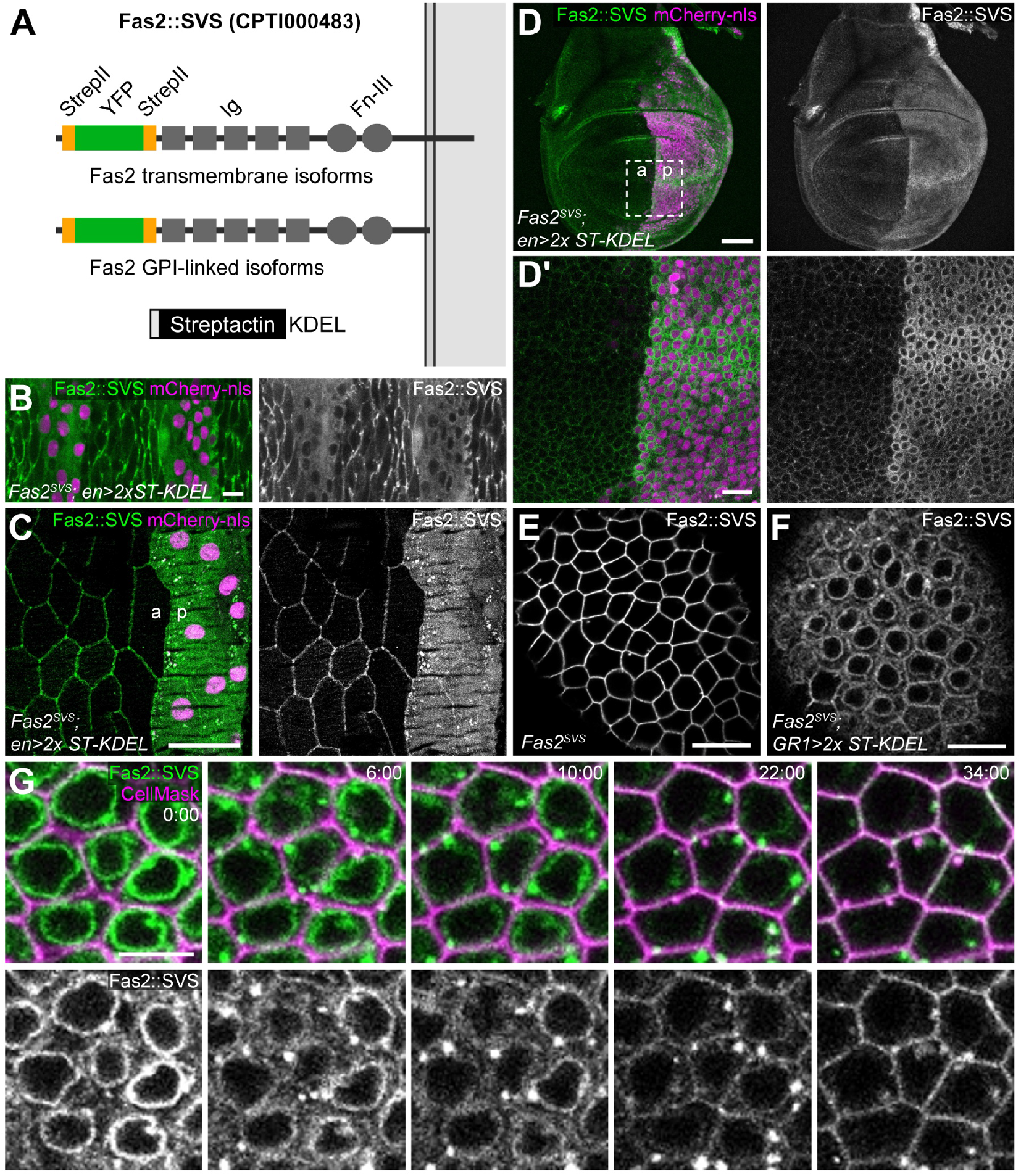
Manipulation of endogenous Fasciclin 2 protein *in vivo* and in cultured ovaries. **(A)** Scheme of Fasciclin 2 (Fas2) protein tagged with venus-YFP (green) flanked by StrepII tags (orange). The *Fas2* locus encodes multiple transmembrane and GPI-linked isoforms, all of which carry the extracellular StrepII-venus-YFP-StrepII (SVS) tag. For simplicity, only a transmembrane isoform is shown. Streptactin-KDEL (bottom) mediates ER retention of secretory proteins carrying StrepII tags in their luminal (extracellular) portion. **(B**,**C)** Maximum-intensity projections of embryonic (B) and third instar-larval (C) epidermis expressing Fas2::SVS (green). Two copies of Streptactin-KDEL and mCherry-nls (magenta) are expressed in epidermal stripes under control of *en-Gal4*. Note that Fas2::SVS outlines lateral membranes of control cells and accumulates in the ER of ST-KDEL-expressing posterior (p) compartment cells marked by mCherry-nls. **(D**,**D’)** Maximum-intensity projection of wing imaginal disc of third-instar larva as in (B,C). Note intracellular retention of Fas2::SVS in posterior (p) compartment. (D’) shows close-up of region marked by square in (D). **(E**,**F)** Confocal section of follicular epithelium in cultured ovarian follicles (stage 7) from *Fas2::SVS* females. Fas2::SVS localizes at lateral membranes in control egg chamber (E), but is retained in the ER in Streptactin-KDEL-expressing follicle cells (F). **(G)** Stills (confocal sections) from time-lapse movie of cultured egg chamber (stage 6) expressing Fas2::SVS (green in G) and Streptactin-KDEL (two copies) in follicle cells. CellMask (magenta) marks plasma membranes. After addition of biotin at t=0 min (1.5 mM final concentration) to the medium, Fas2::SVS is released from the ER, accumulates in puncta (6 min) and becomes detectable at the plasma membrane (22 min). Time (min:s) is indicated. See Movie S8. Scale bars: (B), 10 μm; (C,D), 50 μm; (D’,E,F), 10 μm; (G), 5 μm.

In all cases tested, biotin injection triggered rapid release of the retained proteins from the ER. For instance, the SJ protein Nrx4::SVS was retained in the ER in ST-KDEL-expressing epidermal cells, while it was localized at lateral membranes in adjacent control cells (Fig. S4K). Within four minutes after biotin injection, Nrx4::SVS accumulated in puncta resembling ERES/Golgi clusters. After fifteen minutes, Nrx4::SVS became detectable at lateral plasma membranes (Fig. S4K), suggesting that the newly released Nrx4::SVS protein was incorporated into SJs. Taken together, these findings show that RUSH enables control over ER exit and trafficking of exogenous as well as endogenous transmembrane, GPI-anchored, and secreted proteins in embryos.

### Control of membrane trafficking in organs cultured *ex vivo*

We next tested whether the system can be used also to control secretory trafficking in organs cultured *ex vivo*. To this aim, we cultivated ovarian follicles of females expressing endogenous Fas2::SVS (CPTI 000483; Lye et al., 2014). The NCAM homolog *Fas2* encodes transmembrane and GPI-anchored isoforms (Neuert et al., 2020), all of which carry the SVS tag in the Fas2 ectodomain (Figure 5A). Fas2::SVS localizes at lateral membranes of follicle cells in pre-vitellogenic egg chambers (Fig. 5E). Expression of ST-KDEL in the follicle epithelium of heterozygous *Fas2::SVS/+* females led to retention of Fas2::SVS in the perinuclear ER (Fig. 5F, Fig. S5). Addition of biotin (1.5 mM final concentration) to the culture medium triggered release of Fas2::SVS from the ER after approximately two minutes (n=8; Movie S8). At the same time, Fas2::SVS began to accumulate in large puncta resembling ER exit sites, and became detectable at lateral cell membranes from approximately 20 minutes after biotin addition (Fig. 5G, Movie S8).

### Characterizing the secretory route of Fasciclin2 protein

To visualize intermediate steps of Fas2::SVS trafficking to the plasma membrane, we fixed follicles at different time points after biotin-induced ER-release and labeled specific compartments of the secretory pathway by immunostaining (Fig. S5A-G). In stage 6 follicles, Fas2::SVS was retained in RFP-KDEL-positive perinuclear ER and in ER structures that were enriched in the basal cytoplasm (Fig. S5A). Within ten minutes after biotin addition, Fas2::SVS redistributed to large puncta (0.349 ± 0.1 μm diameter, mean ± s.d.; n=1827 puncta in 286 cells) that localized adjacent to RFP-KDEL-positive ER structures and were labeled by the trans-Golgi marker Golgin245 (Fig. S5B,D), indicating that Fas2::SVS has passed the ER and reached the trans-Golgi network. After 30 minutes, Fas2::SVS appeared in numerous smaller puncta (0.052 ± 0.02 μm diameter, mean ± s.d.; n=786 puncta in 139 cells), which did not overlap with the late-endosomal marker Rab7 (Fig. S5C,E). These Fas2::SVS-positive puncta were distributed along parallel F-actin bundles underneath the basal plasma membrane (Fig. S5G), suggesting that Fas2::SVS-carrying vesicles are transported along basal actin filaments. Indeed, live imaging of follicles 30 minutes after ER release revealed that Fas2::SVS vesicles moved directionally towards the lateral plasma membrane (Fig. S5H,I). The vesicles travelled in parallel to the orientation of the basal F-actin bundles at a speed (0,29 ± 0,1 μm/sec, n=10 vesicles) that is consistent with myosin-based transport (Mehta et al., 1999). Thus, RUSH-based analysis suggests that Fas2 protein is transported through basal ER and Golgi compartments to dynamic vesicles that move, presumably along basal actin filaments, to the plasma membrane. These results demonstrate that RUSH allows to characterize the route of a secretory protein from the ER through the Golgi apparatus to late steps, including transport in secretory vesicles.

### Real-time analysis of tricellular junction assembly

Building on these results, we employed RUSH to investigate the assembly of tricellular junctions (TCJs) in embryonic epithelia. The transmembrane proteins Anakonda (Aka), Gliotactin (Gli) and M6 accumulate at epithelial cell vertices, where they organize TCJs essential for epithelial barrier function, but how TCJ proteins are targeted to vertices is not clear. Analyzing the spatiotemporal dynamics of TCJ assembly has not been possible thus far, because recruitment of TCJ proteins to cell vertices cannot be visualized under steady-state conditions. We therefore used RUSH to synchronize trafficking of the TCJ protein Gli in the embryonic epidermis. Flies carrying the homozygous viable *Gli*^*CPTI002805*^ allele (Lye et al., 2014) produce functional Gli::SVS protein that localizes to TCJs like wild-type Gli protein (Fig. 6A; Wittek et al., 2020). Expression of ST-KDEL in epidermal stripes of heterozygous *Gli*^*CPTI002805*^ embryos led to ER retention of Gli::SVS, while Gli::SVS localized to TCJs in adjacent control cells (Fig. 6A,B). The uniform Gli::SVS signal in the ER of ST-KDEL-expressing cells decreased shortly after biotin injection, and after 2 minutes Gli::SVS accumulated at puncta resembling ERES-Golgi clusters (Fig. 6B,C, Movie S7). Approximately 10 minutes after biotin injection, Gli::SVS signals began to increase sharply at TCJs and to a lesser degree at bicellular junctions (BCJs). Signals at BCJs no longer increased after 15 minutes, while signals at TCJs continued to increase (Fig. 6B,C). Of note, Gli::SVS signal was still visible at BJCs of ST-KDEL-expressing cells 30 minutes after release, whereas Gli::SVS was not visible at BCJs of control cells (Fig. 6B, 30 min). Together, these findings suggest that a fraction of Gli::SVS protein is initially delivered to BCJs and subsequently redistributed to TCJs. Interestingly, the newly released Gli::SVS protein was not incorporated along the entire length of the vertices but accumulated in a single spot at the apical side of each TCJ (Fig. 6D,E, Movie S7). This suggests that growing TCJs are extended in a polarized fashion by addition of new material to the apical side of the junctional complex, resembling the behavior of growing bicellular SJs (Babatz et al., 2018). These findings demonstrate how synchronization of membrane trafficking can be employed to gain insights into the spatiotemporal dynamics of junctional assembly in developing tissues *in vivo*.

**Figure 6.**
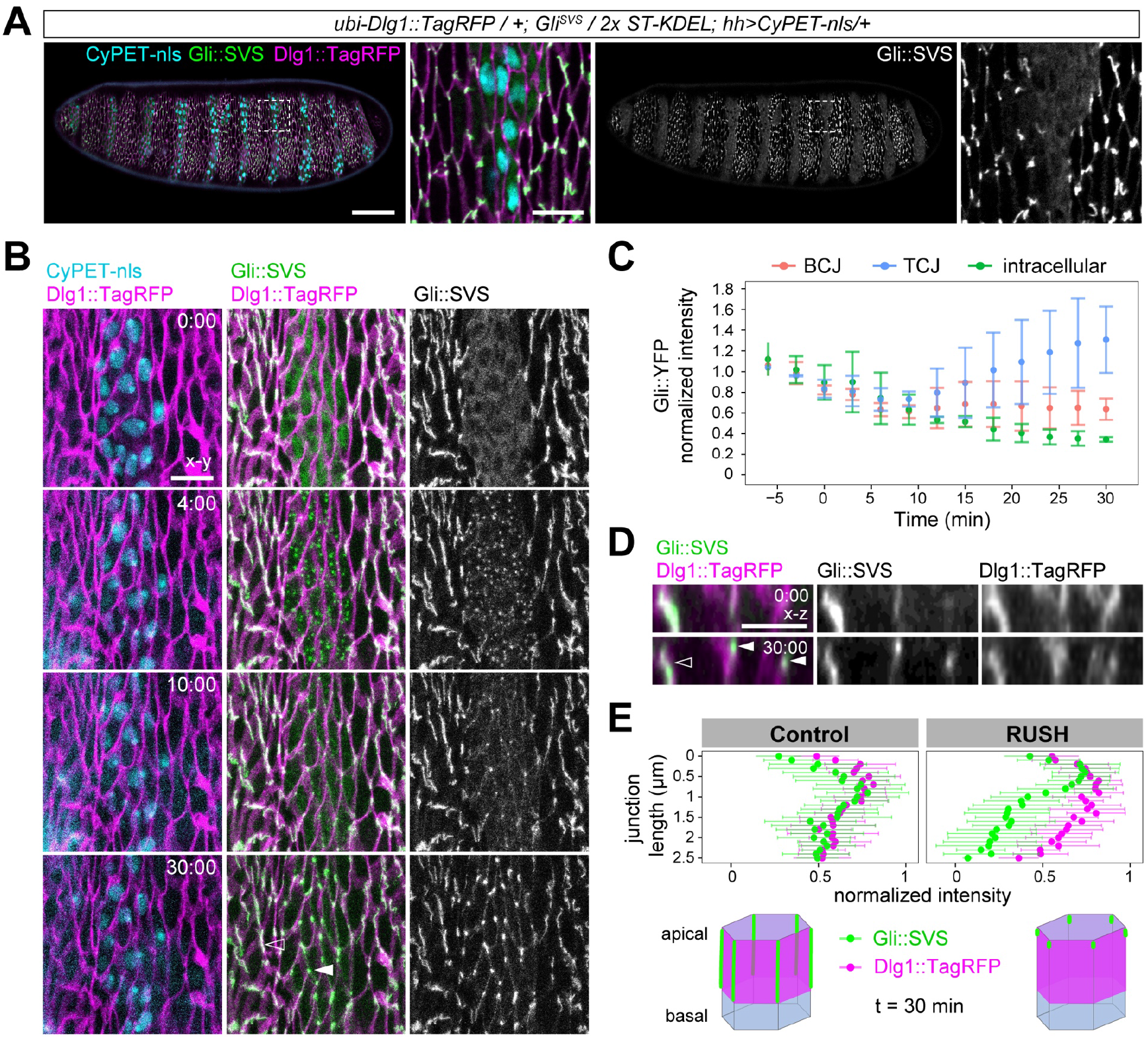
Real-time analysis of tricellular junction assembly in the embryonic epidermis. **(A)** Maximum-intensity projection of living embryo (stage 15) expressing endogenous Gli::SVS (green) and *ubi-*Dlg1::Tag-RFP (magenta). CyPET-nls (cyan) and ST-KDEL (two copies) are expressed in epidermal stripes under the control of *hh-Gal4*. Gli::SVS is retained in the ER of CyPET-nls-marked ST-KDEL-expressing cells and accumulates along tricellular junctions in control cells. Close-ups show region marked by box in overview panels. **(B)** Stills (maximum-intensity projections) from time-lapse movie of embryo as in (A) injected with biotin at t=0 min. After release from ER, Gli::SVS appears at ERES/Golgi puncta (t=4 min) and subsequently begins to accumulate at tricellular junctions (t=10 min). White arrowhead marks newly assembled Gli::SVS at apical tip of vertex in ST-KDEL expressing cell. Open arrowhead marks extended Gli::SVS signal along vertex in control cell. Time (min:s) is indicated. See Movie S7. **(C)** Quantification of Gli::SVS signals at bicellular junctions (BJC, red), tricellular junctions (TCJ, blue) and in the cytoplasm (green). Mean ± s.d. is shown. Per embryo and timepoint, 5 BCJs, 5 TCJs, and 5 cytoplasmic measurements were analyzed in a total of 3 embryos. **(D)** Sections acquired orthogonal (x-z) to the plane of the epithelium at the boundary of ST-KDEL-expressing cells and control cells. Note that newly released Gli::SVS is distributed along the apical-basal length of vertices in control cells (open arrowhead) but accumulates apically at vertices in ST-KDEL-expressing cells (white arrowheads). **(E)** Quantification of Gli::SVS (green) and Dlg1::Tag-RFP (magenta) signals (normalized to maximal signal) along TCJs in control cells (left graph) and ST-KDEL-expressing cells (RUSH; right graph) at t=30 min. TCJs extend from 0 μm (apical) to 2.5 μm (basal). Mean ± s.d. is shown. 5 junctions in control cells and 5 junctions in ST-KDEL-expressing cells per embryo were analyzed in 3 embryos. Scale bars: (A), 50 μm, close-ups: 10 μm; (B), 10 μm; (D), 5 μm.

### Hook-induced ER retention enables tissue-specific interference with protein secretion

Finally, we asked whether hook-induced ER retention could be employed to interfere with secretory protein function. To this aim, we analyzed the effect of ER retention of the Notch (N) receptor on cell fate specification. In the embryonic tracheal system, the N ligand Delta (Dl) is expressed at elevated levels in tracheal fusion cells (FCs) at branch tips and activates N signaling in adjacent stalk cells to prevent these cells from adopting tip cell fate (Fig. 7A). Accordingly, *N* mutations cause excessive FC specification (Ikeya and Hayashi, 1999; Llimargas, 1999; Steneberg et al., 1999). To test whether ST-KDEL-induced ER retention of N protein in tracheal cells can reproduce the tracheal phenotype of *N* mutations, we used the *N*^*SVS-CPTI002347*^ allele (referred to as *N::SVS*), which produces N::SVS protein with an SVS tag inserted into the N ectodomain (Lye et al., 2014). N::SVS protein was weakly detectable in embryonic tracheal (Fig. 7B) and larval wing imaginal disc cells (Fig. 7D). Homozygous (female) and hemizygous (male) *N::SVS* flies were phenotypically normal and fertile, indicating that N::SVS protein is functional. The majority of embryos carrying *N::SVS* showed normal tracheal cell specification with a pair of FCs (discernible by expression of the transcription factor Dysfusion (Dysf)) at each tracheal metamere boundary (Fig. 7B), while supernumerary FCs were found in 12% (n=17) of *N::SVS/+* heterozygous and in 43% (n=14) of *N::SVS/Y* hemizygous embryos (at least one extra FC per embryo; Fig. S6A,B,G). Expression of two copies of ST-KDEL in tracheal cells of *N::SVS* embryos led to intracellular accumulation of N::SVS protein (Fig. 7C) and to excessive specification of FCs in 31% (n=13) of *N::SVS/+* heterozygous and in 76% (n=21) of *N::SVS/Y* hemizygous embryos (Fig. S6C,D,G). Increasing the dosage of *btl*-Gal4 from one to two copies led to fully penetrant (100%; n=11) ectopic FC specification in *N::SVS/Y* hemizygous embryos (Fig. S6F,G) resembling the tracheal defects in N mutants (Ikeya and Hayashi, 1999; Llimargas, 1999; Steneberg et al., 1999), indicating that hook-induced ER retention of N protein effectively blocks N signaling. Moreover, ST-KDEL expression under the control of *en*-Gal4 led to ER retention of N::SVS protein in the posterior compartment of wing imaginal discs in third-instar larvae (Fig. 7D). Adult flies showed characteristic wing margin defects and thickened wing veins, resembling the wing defects of *N*^*1*^ mutants, but restricted to the posterior wing compartment (Fig. 7E-G). Together, these findings demonstrate that hook-induced ER retention can be used to deplete the functional pool of a membrane protein by interfering with its secretion in a tissue-specific manner.

**Figure 7.**
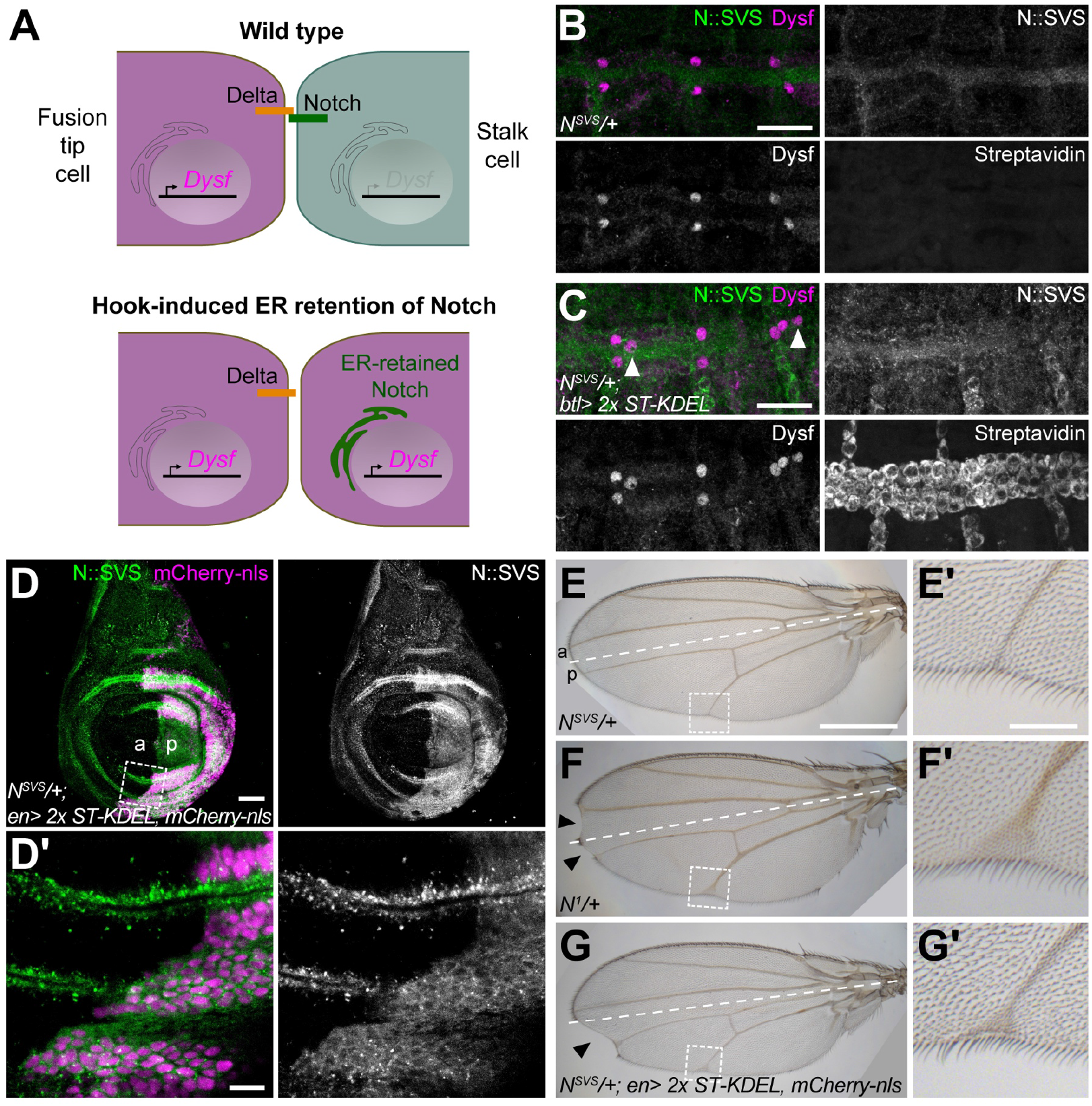
ER retention of Notch::SVS protein causes cell fate specification defects resembling *Notch* loss-of-function mutant. **(A)** Role of Notch signaling in tracheal tip cell specification. The Notch ligand Delta on “fusion” tip cell (magenta) stimulates Notch signaling in adjacent stalk cell (green), preventing it from adopting tip cell fate. ER retention of Notch abolishes Notch signaling and leads to misspecification of stalk cells into tip cells. **(B**,**C)** Representative images of stage 15 *N::SVS/+* control embryo (B) and *N::SVS/+* embryo expressing ST-KDEL (two copies) in tracheal cells (C) stained with anti-GFP (to detect N::SVS; green), anti-Dysfusion (Dysf; magenta) and anti-streptavidin antibodies. Note weak N::SVS signals and a pair of Dysf-positive fusion cells at each tracheal metamere boundary in control embryo (B), whereas *btl*>ST-KDEL-expressing embryo shows intracellular retention of N::SVS and supernumerary Dysf-positive fusion cell nuclei (arrowheads in C). **(D**,**D’)** Live wing imaginal disc of *N::SVS/+* third-instar larva expressing ST-KDEL (two copies) and mCherry-nls in posterior compartment under control of *en-Gal4*. Note retention of N::SVS in the ER of posterior compartment cells marked by mCherry-nls. (D’) shows close-up of boxed region at compartment boundary in (D). **(E-G’)** Wings of control (*N::SVS/+*; E), *N*^*1*^/+ (F), and *N::SVS/+* flies expressing ST-KDEL (two copies) under control of *en-Gal4* (G). Note that ER-retention of N::SVS in the posterior compartment leads to wing margin notches (arrowheads in F,G) and enlarged wing veins resembling the phenotype of *N*^*1*^/+ mutant but restricted to the posterior wing compartment. Close-ups (E’,F’,G’) show enlarged view of distal L5 vein (boxed regions in E-G). Anterior (a) and posterior (p) compartments and the compartment boundary (dashed line) are indicated. Anterior is up. Scale bars: (B,C), 20 μm; (D), 50 μm; (D’), 10 μm; (E,F,G), 500 μm; (E’,F’,G’), 100 μm.

## Discussion

We present a system that enables visualizing the dynamics of membrane trafficking through reversible retention of proteins in the secretory apparatus in a multicellular organism. We show that secreted, transmembrane, and GPI-linked proteins are efficiently retained in the ER by streptavidin hook proteins in *Drosophila* embryos, larvae, and adults. Hook-mediated ER retention is rapidly reversed upon addition of biotin, allowing to synchronize intracellular trafficking and to release fluorescent-marked cargo proteins with high temporal precision in a burst of secretion. The system enables control of exogenous and endogenous proteins, is readily combined with available endogenously tagged protein-trap alleles, and is applicable to various tissues in intact animals, as well to organs cultured *ex vivo*. We demonstrate the utility of this approach for analyzing the kinetics of ER-Golgi-trafficking and protein secretion, and the spatiotemporal dynamics of tricellular junction assembly in embryonic epithelia. Finally, we show that hook-induced ER retention can be employed to deplete the functional pool of proteins, allowing to generate tissue-specific loss-of-function conditions.

While methods for synchronizing secretory trafficking have been used extensively in cultured cells (Boncompain et al., 2012; Chen et al., 2013; Presley et al., 1997; Scales et al., 1997; Weigel et al., 2021), adopting these systems to multicellular organisms has been challenging. A first example of synchronized trafficking in *Drosophila* utilized a regulatable secretory protein (ESCargo, Erv29/Surf4-dependent secretory cargo; Casler et al., 2020) to analyze the kinetics of secretion in a multicellular tissue *ex vivo*. However, this system is based on reversible aggregation of an engineered protein and cannot readily be applied to other cargoes. RUSH represents a versatile two-component system based on short peptide tags (SBP or StrepII) that can be fused to any protein of interest and enable control at physiological temperature by biotin, an endogenous vitamin (Boncompain et al., 2012). Importantly, RUSH employs the physiological KDEL-receptor-based protein retrieval mechanism to achieve ER retention of secretory cargo. This mechanism of protein retention is active in all cells under physiological conditions, and contrasts with other, more artificial experimental systems, which are based on reversible protein aggregation (Casler et al., 2020; Chen et al., 2013; Rollins et al., 2000) or misfolding (Kreis and Lodish, 1986), conditions known to induce the unfolded protein response (Hetz et al., 2020). Secretory cargo is released by dissociating ER-retained streptavidin-cargo complexes using a natural, non-toxic agent. Injection of biotin at concentrations exceeding physiological levels did not cause evident toxicity in *Drosophila* embryos. However, endogenous biotin may reduce the efficacy of hook-induced cargo retention by occupying SBP binding sites on streptavidin. We show that this problem can be overcome by increasing the dosage of streptavidin-KDEL hook proteins. Fly strains carrying multiple copies of streptavidin-KDEL transgenes (Table S1) enable straightforward and versatile application of these tools in genetic crosses.

The use of RUSH revealed rapid ER exit and passage of the Golgi apparatus by secretory proteins in epithelia *in vivo*. While mammalian cells typically contain a single pile of interconnected Golgi stacks with tubular ERES extending between ER and Golgi (Weigel et al., 2021), *Drosophila* cells contain several dispersed Golgi mini-stacks (Kondylis and Rabouille, 2009) adjacent to C- or ring-shaped ERES (Reynolds et al., 2019; Yang et al., 2021). Despite these differences, ER-Golgi trafficking in *Drosophila* tracheal cells occurred on similar time scales (2 to 3 min for Serp-SBG) as described for various GPI-linked and transmembrane RUSH cargos in mammalian cells (Weigel et al., 2021). Overexpressed Serp-SBG was released from the ER with similar kinetics as endogenous proteins (Fas2::SVS, Gli::SVS; Nrx4::SVS), suggesting that rapid ER-Golgi traffic is not due to cargo overexpression, which could in principle activate autoregulatory mechanisms that control secretory flux in response to elevated cargo load (Subramanian et al., 2019). Unlike in mammalian cells, where trafficking of TNF-alpha led to a two-fold size increase of ERES (Weigel et al., 2021), we did not detect obvious changes in the size of mCherry-Sec24-labelled ERES upon ER-release of cargo in tracheal cells. Previous work indicated that subsets of Golgi units in *Drosophila* epithelial cells contain distinct sets of glycosylation enzymes, suggesting possible functional differences between Golgi units (Yano et al., 2005). For the secretory proteins tested here, we did not detect evident differences in the rate of cargo passage between different ERES, all of which appeared to be traversed in an approximately uniform fashion. This suggests that functional differences between individual ER-Golgi units are not reflected by the dynamics of cargo passage through these units.

Our real-time analysis of tricellular junction assembly demonstrates the potential of the RUSH assay for elucidating the intracellular routes of proteins and the kinetics of their assembly into complex structures at the tissue level in living organisms. Importantly, the ability to synchronize and follow the dynamics of these processes in real time reveals insights that are not apparent at steady state or by endpoint analyses, as highlighted by our analysis of tricellular junction assembly. In addition to visualizing secretory trafficking in intact living animals, we demonstrate that hook-induced protein re-localization can be used to interfere with protein function. Manipulating proteins rather than gene expression is required in situations where protein perdurance due to slow turnover upon genetic knockout or RNAi-mediated knock-down precludes loss-of-function phenotypes. While nanobody-based approaches allow to re-localize (Harmansa et al., 2017), trap (Matsuda et al., 2021) or degrade proteins (Caussinus et al., 2012), reversible retention by streptavidin hooks offers the additional potential to acutely restore protein localization and function upon addition of biotin, enabling powerful functional experiments with precise temporal control. For instance, releasing a signaling molecule from a defined source at a defined time will allow to determine the signaling molecule’s mode of dispersal in the tissue and the kinetics of cellular responses across a field of cells.

## Limitations

While RUSH is in principle applicable to any secretory protein, the system requires a short peptide tag (SBP: 38 aa; Keefe et al., 2001; StrepII: 10 aa; Voss and Skerra, 1997) to be inserted into the luminal portion of the protein, without interfering with protein folding or trafficking. We showed that this can be readily achieved for both exogenous and endogenous proteins. Sampling a collection of StrepII-tagged protein trap lines (Lye et al., 2014) revealed that 6 (67%) out of 9 tested proteins are efficiently retained in the ER by streptavidin-KDEL hooks. Approximately 100 such protein trap insertions in membrane-associated or secreted proteins are available (https://kyotofly.kit.jp/stocks/documents/CPTI.html) and provide a valuable resource as potential cargo proteins for RUSH experiments. However, the StrepII tag must be accessible in the luminal part of the protein for binding to streptavidin, and inaccessibility of the StrepII tag may explain the lack of ER-retention observed with some of the tested CPTI lines. Generating additional StrepII-tagged cargo lines is straightforward using recombinase-mediated cassette exchange (RCME; Venken et al., 2011) or CRISPR-Cas9-based genome editing approaches.

Long-term expression of streptavidin-KDEL may cause ER stress due to increased cargo load. However, we found that streptavidin-KDEL expression and ER-retention of RUSH cargo proteins did not generally induce ER stress, but that only high levels of ER hooks induced ER stress in a cell-type-specific manner. Consistent with our findings, certain cell types (*e*.*g*. secretory epithelia) show constitutive induction of XBP1-GFP reporters in the absence of genetic or pharmacological stressors, suggesting that basal activity of the IRE1/XBP1 ER stress sensing system is part of normal *Drosophila* development (Ryoo et al., 2007; Sone et al., 2013). Elevated secretory load was not accompanied by cell death or developmental or morphological abnormalities. However, prolonged streptavidin expression in certain tissues may cause developmental delays due to depletion of biotin, an essential vitamin. This potential problem can be circumvented by tuning the levels or timing of streptavidin expression, *e*.*g*. using temperature-sensitive Gal80 to conditionally control Gal4 activity (McGuire et al., 2004).

While RUSH currently provides temporal control over membrane trafficking, combining the system with orthogonal opto-chemical tools (*e*.*g*. photoactivatable “caged” biotin; Terai et al., 2011) or with photoactivatable or photoconvertible proteins will enable precise spatial and temporal manipulation of cellular processes. We anticipate that the application and further development of the tools presented here will reveal new insights into the dynamics and functions of secretory trafficking in various model organisms.

## Materials and methods

### Fly husbandry and embryo collection

Flies were reared on standard cornmeal-molasses-yeast food. The efficiency of streptavidin-mediated ER retention was strongly dependent on the type of dietary yeast, presumably due to differences in biotin content. Ovaries and embryos from flies that were fed fresh yeast showed more efficient ER retention compared to flies that were fed yeast paste prepared from dry yeast. For RUSH experiments, adult flies were therefore kept on apple juice agar plates with yeast paste prepared from fresh baker’s yeast (Fala) for two days before collection of embryos or ovaries. To reduce the availability of biotin, yeast paste was supplemented with Avidin (50 ppm; Sigma A9275).

### *Drosophila* strains and genetics

Unless noted otherwise, *Drosophila* stocks are described in FlyBase. Cambridge protein trap insertion (CPTI) lines (Lowe et al., 2014; Lye et al., 2014) were obtained from the DGRC stock center (Kyoto, Japan) for Basigin (Bsg; CPTI 100050), Chitin deacetylase-like 4 (Cda4; CPTI 002501), Echinoid (Ed; CPTI 000616), Fasciclin 2 (Fas2; CPTI 000483), Gliotactin (Gli; CPTI 002805), Lachesin (Lac; CPTI 002601), Neurexin 4 (Nrx4; CPTI 001977), Notch (N; CPTI 002347) and Sidekick (Sdk; CPTI001692). Other fly stocks were UAS-mCherry-Sec24 (this work), UAS-Rab4-mRFP (Bloomington 8505), UAS-GRASP65-GFP (Bloomington 8507), UASp-RFP-KDEL (Bloomington 30910), UAS-mCherry-NLS, UAS-CyPET-nls (Caussinus et al., 2008), UAS-palm-mKate2 (Caviglia et al., 2016), UAS-palm-mNeonGreen (Sauerwald et al., 2017), UAS-SerpCBD-GFP (Luschnig et al., 2006), UAS-Xbp1-GFP (Ryoo et al., 2007), UAS-Hsc70-3 RNAi (Bloomington 80420), *ubi-*Dlg1::TagRFP (Pinheiro et al., 2017), *btl*-GAL4, *en*-GAL4, *GR1*-Gal4, *hh*-GAL4, *CyO Dfd-GMR-nvYFP, TM6b Dfd-GMR-nvYFP* (Le et al., 2006). *TM3 Ser Dfd-GMR-nvYFP* was generated by transposase-mediated mobilization of the *P[Dfd-GMR-nvYFP]* P-element from the *FM7i Dfd-GMR-nvYFP* chromosome (Le et al., 2006) onto a *TM3 Ser* chromosome.

### Transgenic constructs

The following cargo and hook constructs were generated in this work: UAS-Serp-SBG (pUAST-SerpCBD-SBP-GFP, pUASp-SerpCBD-SBP-GFP), *ubi*-Serp-SBG (pWRpUbiqPE-SerpCBD-SBP-GFP), UAS-Serp-SBR (pUAST-Serp(CBD)-SBP-mRFP), UAS-streptavidin-KDEL (pUAST-SA-KDEL, pUASTΔSV40-SA-KDEL), UAS-Streptactin-KDEL (pUASTΔSV40-ST-KDEL).

Serp(CBD)-SBP-GFP (Serp-SBG) was generated as follows: A DNA fragment containing the N-terminal portion of the Serp coding sequence including the signal peptide (aa1-25; Luschnig et al., 2006) and chitin-binding domain (CBD), followed by one copy of streptavidin-binding peptide (SBP; 38 aa; Keefe et al., 2001), was synthesized and cloned into pUC57-KanR (GenScript Inc.). The insert was subcloned as an EcoRI-NotI fragment into pUASTattB-EGFP to introduce a C-terminal EGFP tag. The construct was integrated into the attP-3B (VK00037)/22A3 landing site using PhiC31-mediated site-specific integration (Bischof et al., 2007). The Serp(CBD)-SBP-GFP fragment was also subcloned into pUASp to generate pUASp-SerpCBD-SBP-GFP, which was transformed into *y w* flies via P-element mediated transgenesis. For ubiquitous expression under the control of the *ubiquitin* promoter, the Serp-SBG coding sequence was inserted into pWRpUbiqPE and transformed into *y w* flies via P-element mediated transgenesis.

Serp(CBD)-SBP-mRFP (Serp-SBR) was generated as follows: A DNA fragment comprising the N-terminal portion of the Serp coding sequence, an SBP tag and the mRFP coding sequence was synthesized and cloned into pUC57-KanR (GenScript Inc.) and subcloned as an EcoRI-XBaI fragment into pUASTattB (Bischof et al., 2007). The construct was integrated into the attP40/25C6 landing site.

Streptavidin-KDEL (SA-KDEL) was generated by fusing the core SA coding sequence (UniProt P22629; aa 36-163; reverse-translated using the *Drosophila melanogaster* codon distribution) to the N-terminal signal peptide of Serp (aa 1-25) and to a C-terminal four-amino-acid ER retention signal (KDEL). The SA-KDEL protein sequence was reverse-translated using the *Drosophila melanogaster* codon usage distribution, synthesized, cloned into pUC57-KanR (GenScript, Inc.), and subcloned as an EcoRI-XbaI fragment into pUASTattB (Bischof et al., 2007) and pUASTΔSV40attB, respectively. pUASTΔSV40attB is a pUASTattB derivative lacking the SV40 3’-UTR that targets transcripts for nonsense-mediated mRNA decay. Deletion of the intron-containing 3’-UTR results in stabilized transcripts and approximately 5-fold higher expression levels compared to pUAST transgenes (Nelson et al., 2018). The pUASTΔSV40attB-SA-KDEL construct was integrated into the attP40/25C6, attPZH-51C1, attP-3B(VK00031)/62E1 and attPZH-86Fa landing sites. The pUAST-SA-KDEL construct was integrated into the attP2/68A4 landing site.

Streptactin-KDEL (ST-KDEL) was generated by fusing the ST coding sequence (Voss and Skerra, 1997) to the N-terminal signal peptide of Serp (aa 1-25) and a C-terminal ER retention signal (KDEL). This fragment was synthesized, inserted in pUC57-KanR (GenScript, Inc.), and subcloned as an EcoRI-KpnI fragment in pUASTΔSV40attB. The construct was integrated into the attPZH-2A, attP18/6C12, attP4/12C6 (X-chromosome), attP40/25C6, attPZH-51C1 (second chromosome), attP-3B(VK00031)/62E1 and attPZH-86Fa (third chromosome) landing sites.

To increase the copy number of UAS-SA-KDEL or UAS-ST-KDEL transgenes, two insertions on the X-chromosome (attPZH-2A, attP18/6C12), second chromosome (attP40/25C6, attPZH-51C1), or third chromosome (attP-3B(VK00031)/62E1, attPZH-86Fa), respectively, were recombined. A strain carrying four copies of UAS-SA-KDEL was generated by combining two insertions on the second chromosome (attP40/25C6, attPZH-51C1) with two insertions on the third chromosome (attP-3B(VK00031)/62E1, attPZH-86Fa).

mCherry-Sec24 was generated by amplifying the *Sec24AB* coding region (including introns) from genomic DNA of Oregon R flies using oligonucleotides BamHI-sec24-F (5’-ATATGGATCCATGTCGACTTACAATCCGAAC) and sec24-XbaI-R (5’-TATATCTAGATCGCTTCGTGCTTCTAGTCA). The PCR product was cut with BamHI and XbaI and the resulting fragment was inserted into pUASTattB-mCherry-mcs (BglII-XbaI) to fuse mCherry to the N-terminus of Sec24. The pUASTattB-mCherry-Sec24 construct was integrated into the attP/ZH-86Fa landing site. After establishing a homozygous viable transformant line, the *w*^*+*^ and *3xP3-RFP* markers of the attP/ZH-86Fa landing site were removed using Cre-mediated recombination.

UAS-Fas2B-SVS (GPI-anchored Fas2 isoform B, carrying the SVS tag at the same position as in the *Fas2*^*CPTI000483*^ protein trap allele) was generated by isolating *Fas2* cDNAs from *Fas2*^*CPTI000483*^ homozygous females. Total RNA was extracted from ovaries using TRIzol reagent (Thermo Fisher, 15596026). polyA+ mRNA was isolated using Dynabeads mRNA purification kit (Thermo Fisher, 61006). First-strand cDNA was generated using ProtoScript first strand cDNA synthesis kit (NEB, E6300S) with oligo-dT primers. cDNAs corresponding to the *Fas2-B* isoform was amplified using isoform-specific oligonucleotides for *Fas2-B* (Fas2-F: 5’-TCATACTCGCATTCTCTCGC and Fas2B-R: 5’-TGATAATTTGTCAGCGGGAGG). PCR products were cloned into pCR-BluntII-Topo vector using Zero Blunt Topo PCR Cloning Kit (Thermo Fisher, 450245), sequenced, and subcloned into the EcoRI site in pUAST-attB. The construct was integrated into the attP40/25C6 and attP2/68A4 landing sites.

### Antibodies and immunostainings

Embryos were fixed in 4% formaldehyde in PBS/heptane for 20 minutes and devitellinized by shaking in methanol/heptane. Ovaries were dissected in M3 insect medium (Sigma S8398) and transferred into a 2 ml Eppendorf tube. For RUSH experiments, medium was removed and 400 μl new M3 medium containing 1.5 mM Biotin was added for either 10 or 30 min. After fixation for 10 min in 4% formaldehyde in PBS, ovaries were permeabilized by washing with 0.1% TritonX-100 in PBS (PBT), blocked for 20 min with 0.5% BSA in PBT, and then incubated with primary antibodies for 3 h at room temperature. For detecting F-actin, Alexa Fluor 568-Phalloidin (Thermo Fisher, A12380) was added at 1:100 dilution. After three washing steps, ovaries were incubated for 2 h with secondary antibodies, washed and mounted in Vectashield mouting medium.

The following antibodies were used: chicken anti-GFP (1:500; Abcam 13970), mouse anti-GFP mAB 12A6 (1:300; DSHB), goat anti-GFP-FITC (1:100; GeneTex GTX26662-100), mouse anti-mCherry (1:200; Biorbyt orb66657), rabbit anti-RFP (1:200; Rockland 600-401-379), chicken anti-mCherry (1:1000; Novus NBP2-25158SS), mouse anti-Streptavidin (1:500; Abcam S10D4, ab10020), rabbit anti-Streptavidin (1:200; Abcam ab6676), rabbit anti-Serp (1:300; Luschnig et al., 2006), goat anti-Golgin245 (1:400; DSHB), mouse anti-Rab7 (1:20; DSHB), mouse anti-KDEL (1:100; Abcam 10C3), rabbit anti-Dysfusion (1:500; Jiang and Crews, 2003), rabbit-anti-Sec16 (1:1000; Ivan et al., 2008), mouse anti-Sxl M18-c (1:500; DSHB), anti-Tango (1:200; DSHB). Goat secondary antibodies were conjugated with Alexa Fluor 488, Alexa Fluor 568, Alexa Fluor 647 (Thermo Fisher), or Cy5 (Jackson ImmunoResearch). Chitin was detected using AlexaFluor-labeled chitin-binding domain from *Bacillus circulans* chitinase A1 as described in (Caviglia and Luschnig, 2013).

### Culture of ovarian follicles

Adult females expressing Fas2::SVS (CPTI000483) and two copies of ST-KDEL in follicle cells under the control of *GR1*-Gal4 were fed fresh yeast paste supplemented with Avidin (50 ppm; Sigma A9275) at 27°C for two days after eclosion. Ovaries were dissected in M3 insect medium containing 0.05% KHCO_3_ and 1x penicillin/streptomycin (from 100x stock; Thermo Fisher). To minimize biotin content of the medium, yeast extract and Bacto Peptone were not added. CellMask Orange (Thermo Fisher) and Hoechst 33342 (Sigma) were added to the medium to stain plasma membranes and nuclei, respectively. Dissected follicles were transferred to 8-well glass-bottom chambers (VWR 734-2061) containing 200 μl M3 medium per well. Before use, glass-bottom chambers were coated with poly-D-Lysine (Sigma P1024; 1 mg/ml in water, pH 8.5) for 1 h at 37°C to immobilize follicles on the glass surface. To induce release of ER-retained proteins, 200 μl of M3 medium containing 3 mM biotin were added to the culture medium (1.5 mM final biotin concentration) during imaging. For inducing ER stress, Dithiothreitol (DTT; Roth 6908.2) was added to the culture medium (5 mM final concentration) for 2.5 hours before imaging.

### Microscopy and embryo microinjections

Imaging was performed using HyD detectors and 40x/1.3 NA and 63x/1.4 NA objectives on a Leica SP8 inverted confocal microscope or using 40x/1.3 NA and 60x/1.35 NA objectives on an Olympus FV1000 inverted confocal microscope. For live imaging, embryos (12-15 h AEL) were dechorionated, placed on apple juice agar plates, and visually selected for ER retention of GFP-tagged cargo proteins using a Leica M165 FC fluorescence stereomicroscope. Selected embryos were transferred to glue-coated coverslips and covered with Voltalef 10S halocarbon oil (VWR) before injection with D-biotin (B4501; Sigma-Aldrich) at the indicated concentration (unless indicated otherwise, 1 mM in water). Biotin was injected ventrally into the body cavity of embryos using either a Transjector 5246 microinjector (Eppendorf) with Femtotips II at a transmitted light microscope or using a FemtoJet microinjector (Eppendorf) mounted on an inverted Leica SP8 confocal microscope.

For high-resolution imaging of ER exit sites, living embryos were imaged using a 63x/1.4 NA objective on a Leica SP8 confocal microscope. Images were processed by deconvolution with Huygens Professional (Scientific Volume Imaging, The Netherlands, http://svi.nl) using the CMLE algorithm with SNR:15 and 40 iterations.

For tracking Fas2::SVS vesicles in follicle cells, cultured egg chambers were imaged on an inverted Leica SP5 confocal microscope using HyD detectors and a 63x/1.30 NA HCX PL APO glycerol immersion objective. For time-lapse movies, images (512 × 256 pixels) were acquired at 1 second time intervals.

### Image analysis

Images were processed using Fiji/ImageJ (Schindelin et al., 2012), Imaris (v7.7.0; Bitplane), OMERO (5.4.10; Allan et al., 2012), and Scikit-Image (v0.18.2; van der Walt et al., 2014). Serp-SBG signals in tracheal cells and lumen were measured in manually selected regions of interest (ROIs) on average-intensity projections of confocal sections acquired in tracheal DT metamere 7. Background intensity was measured in the hemocoel outside the trachea and was subtracted from intracellular and luminal signals. Serp-SBG-containing endosomal vesicles were segmented by applying a median filter (radius 2.0) and a manual threshold. Vesicle diameter (Feret distance) was measured using the Analyze Particles plugin in Fiji. Vesicle velocity was measured using the Spots function in Imaris.

For analyzing the dynamics of Serp-SBG at ER exit sites, mCherry-Sec24-labeled ERES were segmented using the Imaris 3D Spot function. Serp-SBG signals within spherical mCherry-Sec24-containing volumes were measured in Imaris. To quantify the area of intersection between mCherry-Sec24 and Serp-SBG signals at ERES, signals were segmented using Default (for mCherry-Sec24) or Yen Fiji (for Serp-SBG) automatic threshold methods.

For analysis of tricellular junction assembly in embryonic epidermis, images were processed by deconvolution and corrected for x-y drift using the CMLE algorithm in Huygens Professional with SNR:10 for the CyPet-nls channel, SNR:8 for Gli::SVS and Dlg1::tagRFP channels, and 30 iterations (all channels). For quantifying Gli::SVS signals, mean gray values at TCJs, BCJs, and inside cells were measured in manually selected ROIs for each time point on average-intensity projections of the three most apical slices showing junctional signal in a z-stack. For each time point, the mean intensity for TCJs and intracellular signals, respectively, was calculated. Values were then normalized to the mean of the first three time points per group (TCJs, BCJs, intracellular signals). To determine the apical-basal distribution of Gli::SVS along vertices, Gli::SVS signals were analyzed in average-intensity projections of three vertical (x-z) sections. Dlg1::tagRFP signals were used to manually trace vertices of control and ST-KDEL-expressing cells. Gli::SVS signals were measured along a line (width = 10 pixels) drawn along the Dlg1::tagRFP-marked vertex. Values for each junction were normalized to the highest intensity value measured for the respective junction.

For quantifying nuclear Xbp1-GFP signals as a read-out of ER stress, nuclei were segmented using Hoechst 33342 staining (for follicle cells in cultured egg chambers) or using nuclear Tango staining (for tracheal cells in fixed embryos). A Median filter (radius = 8) was applied to the nuclear signal, followed by image binarization using a Mean threshold. Small particles were removed by size exclusion. Xbp1-GFP signals were measured within segmented nuclei using FIJI (v2.3.0). Mean Xbp1-GFP intensities per nucleus and mean values of all nuclei per sample were calculated using R. At least 20 nuclei per embryo and at least 8 nuclei per egg chamber were analyzed.

Fas2::SVS vesicles were tracked using the FIJI Manual tracking plug-in. For illustration of individual tracks, the tracks were recolored and then assembled according to their orientation and size in a standardized epithelial cell using Adobe Illustrator.

### Statistics

Statistical analyses were performed in Microsoft Excel 2010, OriginLab 8.5, or R (3.5.1) using RStudio Interface (1.3.1093). Sample size (n) was not predetermined using statistical methods but was assessed empirically by considering the variability of a given effect as determined by the standard deviation. Experiments were considered independent if the specimens analyzed were derived from different crosses. Investigators were not blinded to allocation during experiments. The data was tested for normality using the Shapiro-Wilk test. Student’s t-test was used for normally distributed data. When the data was not normally distributed, the Wilcoxon rank-sum test (R standard package) was used. P values were corrected for multiple testing using the Bonferroni-Holm method (Holm, 1979). Sample size is indicated in the graphs or figure legends.

## Author contributions

Conceptualization, J.G., C.C., S.L.; Methodology, all authors; Investigation, J.G., C.C., M.H., J.I.S., T.J., W.B., N.B., V.R.; Formal Analysis, J.G., C.C., M.H., T.J., R.S., N.B., V.R; Visualization, J.G., C.C., M.H., T.J., W.B., N.B., V.R.; Reagents and tools, J.G., C.C., M.H., T.J., W.B., D.F., S.L.; Writing – Original Draft, J.G., C.C., S.L.; Writing – Review and Editing, C.C., S.L.; Funding Acquisition, V.R., S.L.; Supervision, S.L.

## Acknowledgements

We thank Franck Perez for advice and discussions, and Sarah Weischer and Thomas Zobel at the Münster Imaging Network for expert support with image analysis. N.B. and V.R. acknowledge support by the Core Facility Live Cell Imaging Mannheim (DFG INST 91027/9-1 FUGG).

