## Supplemental Information for "Acute manipulation and real-time visualization of membrane trafficking and exocytosis in *Drosophila*"

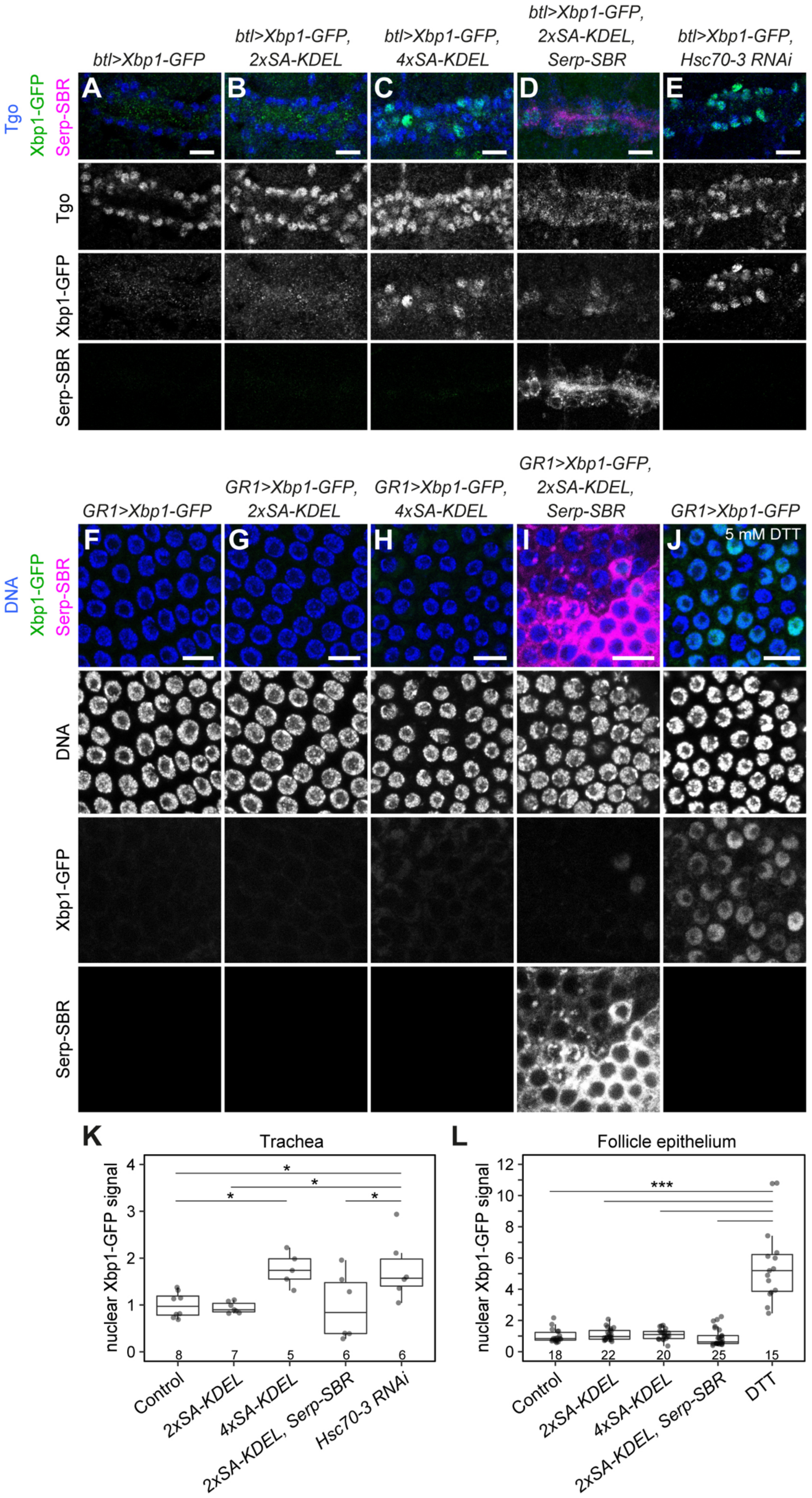
btl>Xbp1-GFP
btl>Xbp1-GFP, 2xSA-KDEL
btl>Xbp1-GFP, 4xSA-KDEL
btl>Xbp1-GFP, 2xSA-KDEL, Serp-SBR
btl>Xbp1-GFP, Hsc70-3 RNAi
Tgo
Xbp1-GFP
Serp-SBR
A
B
C
D
E
GR1>Xbp1-GFP
GR1>Xbp1-GFP, 2xSA-KDEL
GR1>Xbp1-GFP, 4xSA-KDEL
GR1>Xbp1-GFP, 2xSA-KDEL, Serp-SBR
GR1>Xbp1-GFP
5 mM DTT
DNA
F
G
H
I
J
K
Trachea
nuclear Xbp1-GFP signal
Control
2xSA-KDEL
4xSA-KDEL
2xSA-KDEL, Serp-SBR
Hsc70-3 RNAi
L
Follicle epithelium
DTT
***

**Figure S1**

**Streptavidin-KDEL expression induces dosage-dependent ER stress in embryonic tracheal cells, but not in ovarian follicle cells. Related to Figures 1 and 2.**

**(A-E)** Confocal sections of tracheal dorsal trunk in embryos (stage 15) of the indicated genotypes stained with anti-GFP (to detect Xbp1-GFP; green), anti-Tango (Tgo, tracheal cell nuclei; blue) and anti-RFP (to detect Serp-SBR; magenta) antibodies. Note that nuclear Xbp1-GFP signals are not detectable in control embryos expressing only Xbp1-GFP under control of *btl-Gal4* (A) or in embryos co-expressing two copies of SA-KDEL (B), but are detectable in nuclei of a subset of tracheal cells co-expressing four copies of SA-KDEL (C). ER-retention of Serp-SBR (magenta) in SA-KDEL-expressing cells does not lead to elevated nuclear Xbp1-GFP signals (D), whereas *Hsc70-3* RNAi leads to Xbp1-GFP accumulation, indicative of ER stress (E).

**(F-J)** Confocal sections of follicle epithelium in living egg chambers (stage 9) expressing Xbp1-GFP (green) and stained with Hoechst 33342 to detect nuclei (blue). Note that nuclear Xbp1-GFP is not detectable in control follicles (F) and in follicles expressing either two (G) or four (H) copies of SA-KDEL, and is detectable only in a small subset of nuclei in follicles co-expressing SA-KDEL and Serp-SBR (magenta; I). By contrast, follicles incubated with DTT (5 mM) show nuclear accumulation of Xbp1-GFP indicative of ER stress (J).

**(K,L)** Quantification of nuclear Xbp1-GFP signals in embryonic tracheal cells (K) and ovarian follicle cells (L). Values were normalized to the mean of the respective control sample. Each data point represents the mean of at least 20 nuclei analyzed in one embryo (K) or the mean of at least 8 nuclei analyzed in one egg chamber (L). Number of embryos (K) or egg chambers (L) analyzed is indicated for each genotype. Boxplot shows the median (line), interquartile range (box) and 1.5x interquartile range from the 25th and 75th percentile (whiskers). p-values (K, pairwise t-test; L, Wilcoxon rank-sum test) are indicated.

Scale bars: (A-J), 10 µm.

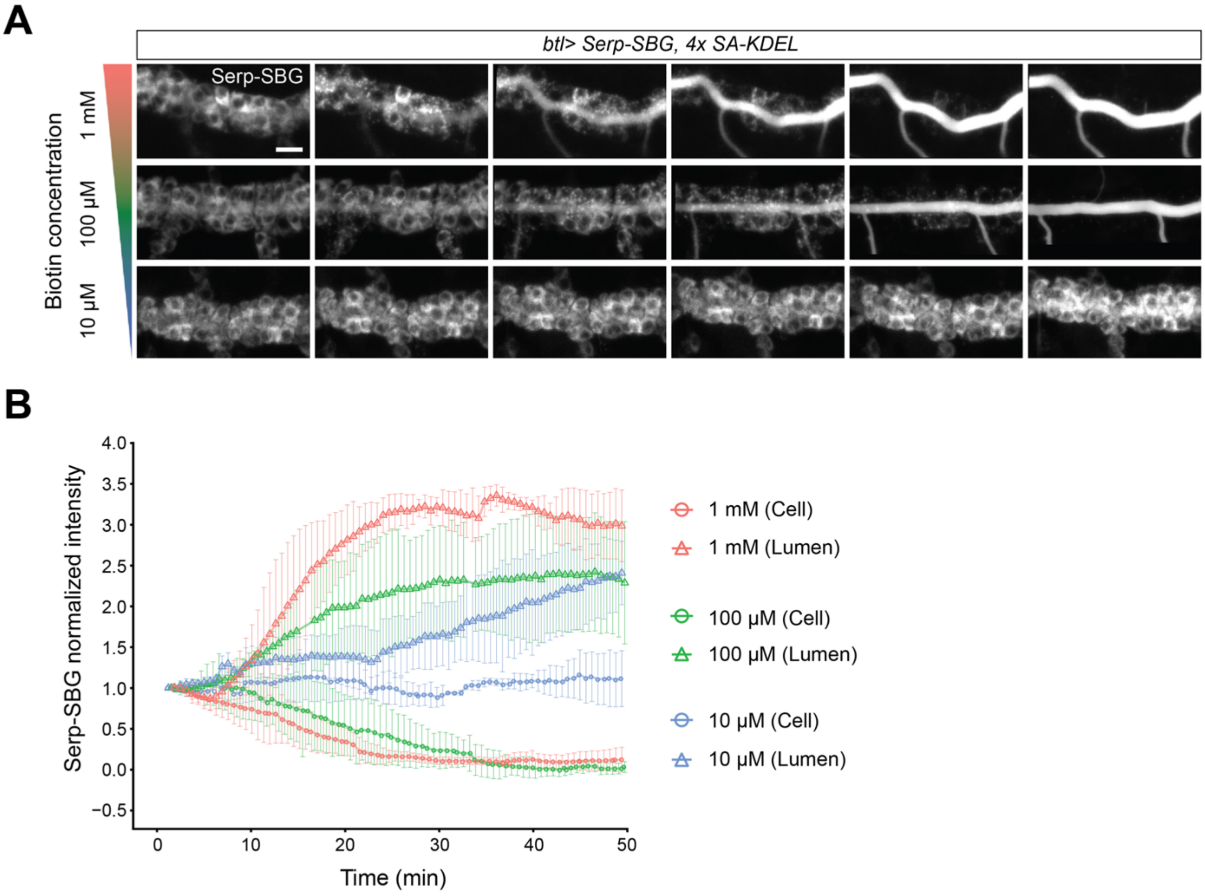


**Figure S2**

**Effect of biotin concentration on ER release of Serp-SBG. Related to Figure 3.**

**(A)** Representative images of embryos expressing Serp-SBG and copies of SA-KDEL in tracheal cells after injection of different concentrations (1 mM, 100 µM or 10 µM) of biotin at t=0 min. Time (min:s) after injection is indicated. See Movie S4.

**(B)** Quantification of Serp-SBG signals in tracheal cells (circles) and lumen (triangles) of embryos injected with 1 mM (red), 100 µM (green) or 10 µM biotin (blue). Values were normalized to intensities at t=0. Mean +/- s.d. is shown. n=4 embryos per concentration.

Scale bar (A): 10 µm.

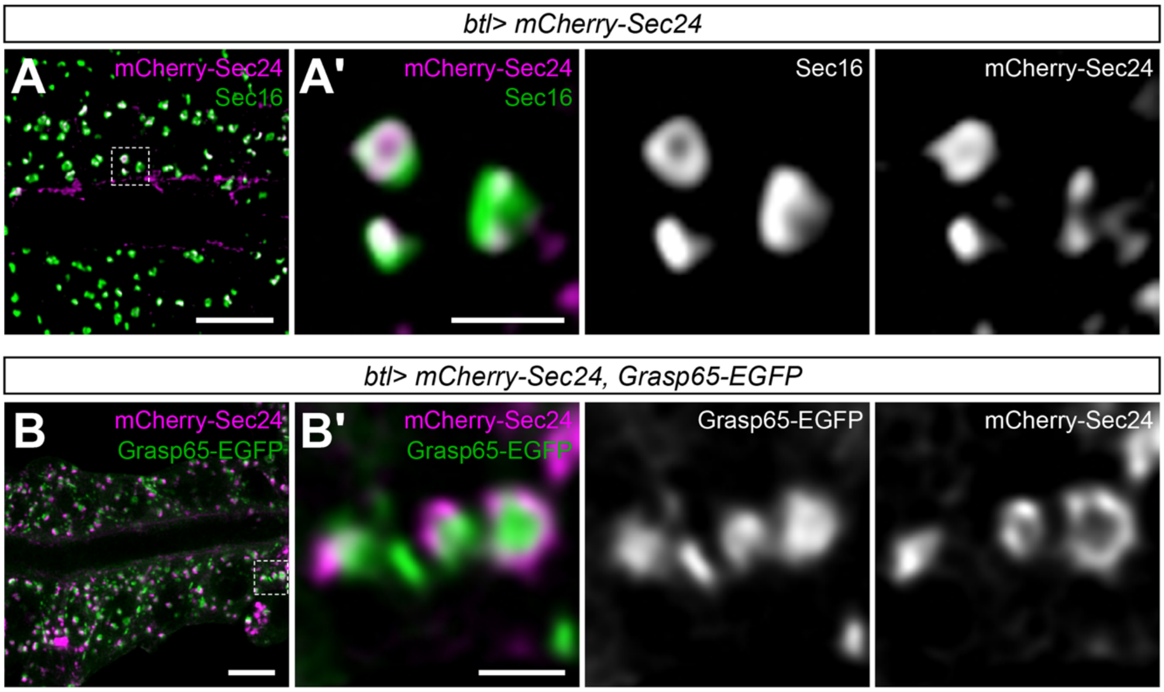


**Figure S3**

**mCherry-Sec24 marks ER exit sites adjacent to Golgi stacks. Related to Figure 4.**

**(A,A')** Confocal maximum-intensity projection (A) and single section (A') of tracheal dorsal trunk cells in embryo (stage 15) expressing mCherry-Sec24 (magenta) and stained for Sec16 (green).

**(B,B')** Confocal maximum-intensity projection (B) and single section (B') of tracheal dorsal trunk cells in embryo (stage 15) expressing mCherry-Sec24 (magenta) and GRASP65-EGFP (green).

Note that mCherry-Sec24 colocalizes with the ERES marker Sec16 (A,A'), but localizes adjacent to GRASP65-EGFP-marked Golgi stacks (B,B'). (A') and (B') show close-ups of the regions marked in (A,B).

Scale bars: (A,B), 5 µm; (A',B'), 1 µm.

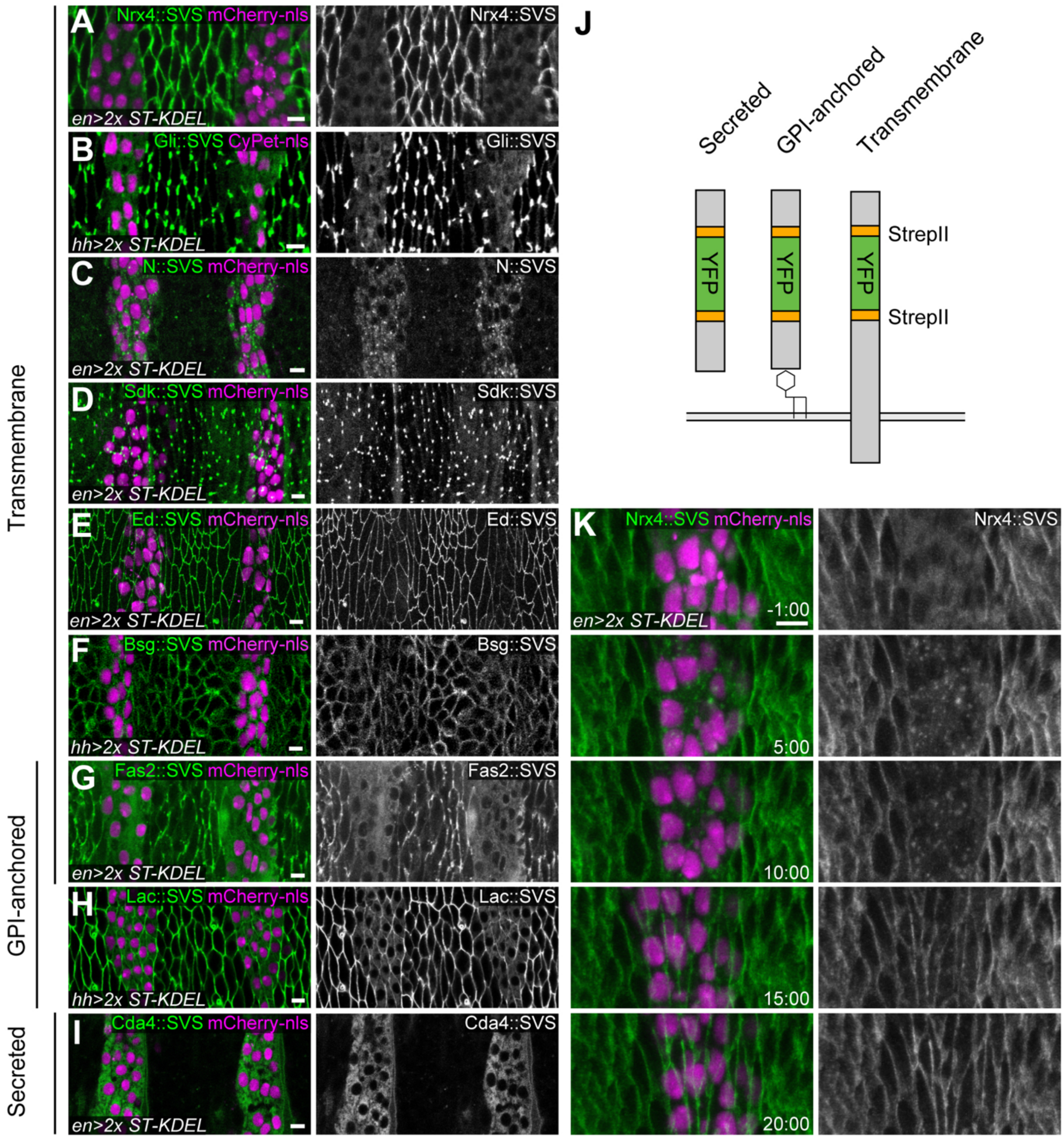


**Figure S4**

**RUSH enables manipulation of StrepII-tagged endogenous proteins. Related to Figure 5.**

**(A-I)** Confocal maximum-intensity projections of lateral epidermis in embryos (stage 15) expressing different SVS-tagged protein traps (green). ST-KDEL (two copies) and mCherry-nls (magenta) are expressed in epidermal stripes under the control of *en*-Gal4 or *hh*-Gal4 as indicated. Representative images of protein traps in transmembrane (Neurexin 4 (Nrx4, CPTI

001977; A), Gliotactin (Gli, CPTI 002805; B), Notch (N, CPTI 002347; C), Sidekick (Sdk, CPTI001692; D), Echinoid (Ed, CPTI 000616; E), Basigin (Bsg, CPTI 100050; F)), GPI-anchored (Fasciclin 2 (Fas2, CPTI 000483; G), Lachesin (Lac, CPTI 002601; H)) and secreted (Chitin deacetylase 4 (Cda4, CPTI 002501; I)) proteins are shown. Note that most SVS-tagged proteins are retained in the ER of ST-KDEL-expressing cells. However, no efficient ER retention was observed for Sdk::SVS (D), Ed::SVS (E), and Bsg::SVS (F). *Fasciclin2* encodes transmembrane and GPI-anchored isoforms, all of which carry the SVS tag in the Fas2::SVS (CPTI 000483) line (G). Cda4::SVS (I) is detectable only in ST-KDEL-expressing cells because the protein is secreted apically (out of the focal plane) by cells not expressing ST-KDEL.

**(J)** Schematic illustration of transmembrane, GPI-anchored, and secreted proteins (grey), tagged with venus-YFP (green) flanked by StrepII tags (orange) in their luminal (extracellular) part.

**(K)** Stills from time-lapse movie of lateral epidermis in Nrx4::SVS embryo expressing ST-KDEL (two copies) and mCherry-nls under the control of *en-gal4*. Biotin was injected at t=0 min. Time (min:s) is indicated. Note that Nrx4::SVS is retained in the ER of ST-KDEL-expressing cells (marked by mCherry-nls). 5 min after biotin injection Nrx4::SVS accumulates at ERES/Golgi stacks and becomes detectable at lateral plasma membranes 15 min after injection.

Scale bars: (A-I,K), 5 µm.

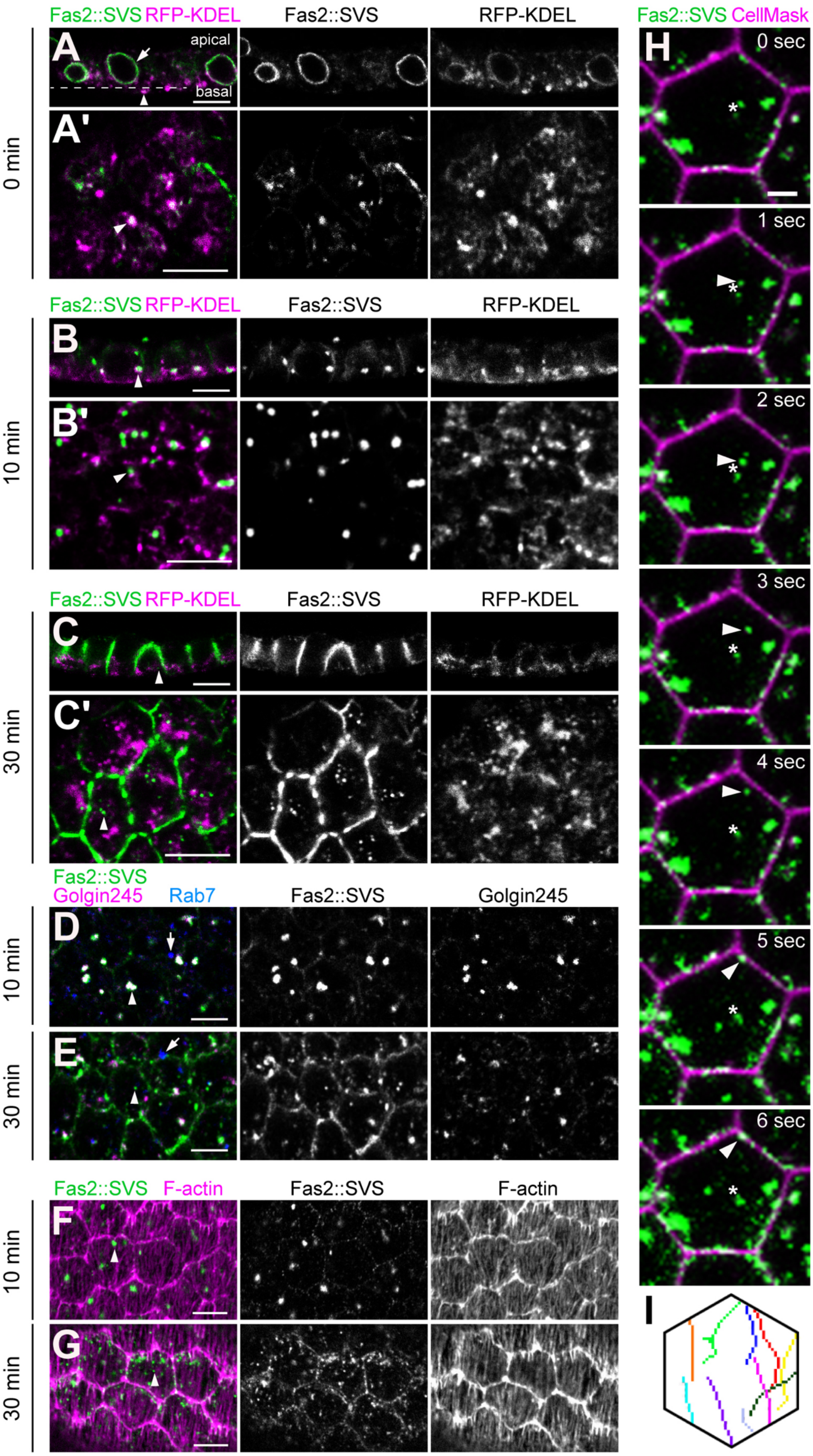
Fas2::SVS RFP-KDEL
Fas2::SVS
RFP-KDEL
A
apical
basal
A'
0 min
B
B'
10 min
C
C'
30 min
Fas2::SVS
Golgin245
Rab7
Golgin245
D
10 min
E
30 min
Fas2::SVS F-actin
F-actin
F
10 min
G
30 min
Fas2::SVS CellMask
H
0 sec
1 sec
2 sec
3 sec
4 sec
5 sec
6 sec
I

**Figure S5**

**RUSH reveals steps of Fas2 trafficking in follicle cells. Related to Figure 5.**

**(A-C’)** Confocal sections of follicular epithelium in cultured egg chambers (stage 6) expressing endogenous Fas2::SVS (green), RFP-KDEL (magenta) and Streptactin-KDEL under control of GR1-Gal4. Follicles were fixed just before (A,A’), 10 minutes after (B,B’), or 30 minutes after (C,C’) addition of biotin, and were immunostained for GFP and RFP. (A,B,C) show sections along the apical-basal axis (x-z plane). (A’,B’,C’) show orthogonal sections in the basal region (dashed line; x-y plane). (A,A’) Before addition of biotin, Fas2::SVS localizes in the perinuclear (arrow) and cytoplasmic (arrowhead) ER, where it colocalizes with RFP-KDEL. (B,B’) 10 minutes after biotin addition, Fas2:SVS accumulates in large puncta adjacent to basally enriched RFP-KDEL-positive ER structures (arrowheads). (C,C’) 30 minutes after biotin addition, Fas2::SVS is detectable at the plasma membrane and in small vesicles (arrowheads) distributed in the basal cytoplasm.

**(D,E)** Basal sections of stage 6 follicles expressing endogenous Fas2::SVS (green) and Streptactin-KDEL under control of GR1-Gal4. Follicles were fixed 10 minutes (D) or 30 minutes (E) after addition of biotin, and were immunostained for GFP (Fas2::SVS, green), Golgin245 (magenta) and Rab7 (blue). (D) 10 minutes after biotin addition, Fas2::SVS colocalizes with Golgin245-positive trans-Golgi compartments (arrowhead) but not with Rab7-positive endosomes (arrow). (E) 30 minutes after biotin addition, Fas2::SVS is detectable at the plasma membrane and in small vesicles (arrowhead) that do not overlap with Golgi compartments or Rab7-positive endosomes (arrow).

**(F,G)** Basal sections of stage 6 follicles expressing endogenous Fas2::SVS (green) and Streptactin-KDEL under control of GR1-Gal4. Follicles were fixed 10 minutes (F) or 30 minutes (G) after addition of biotin, and were stained for GFP (Fas2::SVS, green) and with phalloidin to visualize F-actin (magenta). (F) 10 minutes after biotin addition, a small number of Fas2::SVS puncta (arrowhead) resembling Golgi compartments (D) are detected. (G) 30 minutes after biotin addition, many small Fas2::SVS-containing vesicles (arrowhead) are detected near basal F-actin bundles.

**(H)** Stills from representative time-lapse movie of follicle (stage 6, basal section) expressing UAS-Fas2::SVS (green) and Streptactin-KDEL under control of GR1-Gal4, 30 minutes after biotin addition. CellMask (magenta) marks plasma membranes. The arrowhead indicates a Fas2::SVS-containing vesicle that moves from the central region to the plasma membrane. The starting point of the vesicle track is marked by an asterisk. Time is indicated.

**(I)** Tracks of 10 independently recorded Fas2::SVS-positive vesicles that moved from the cell interior to the lateral plasma membrane. Tracks were assembled according to their orientation and size in a standardized cell. Note that vesicle tracks are oriented parallel to the basal F-actin bundles (F,G).

Scale bars: (A-G), 5 μm, (H), 1 μm.

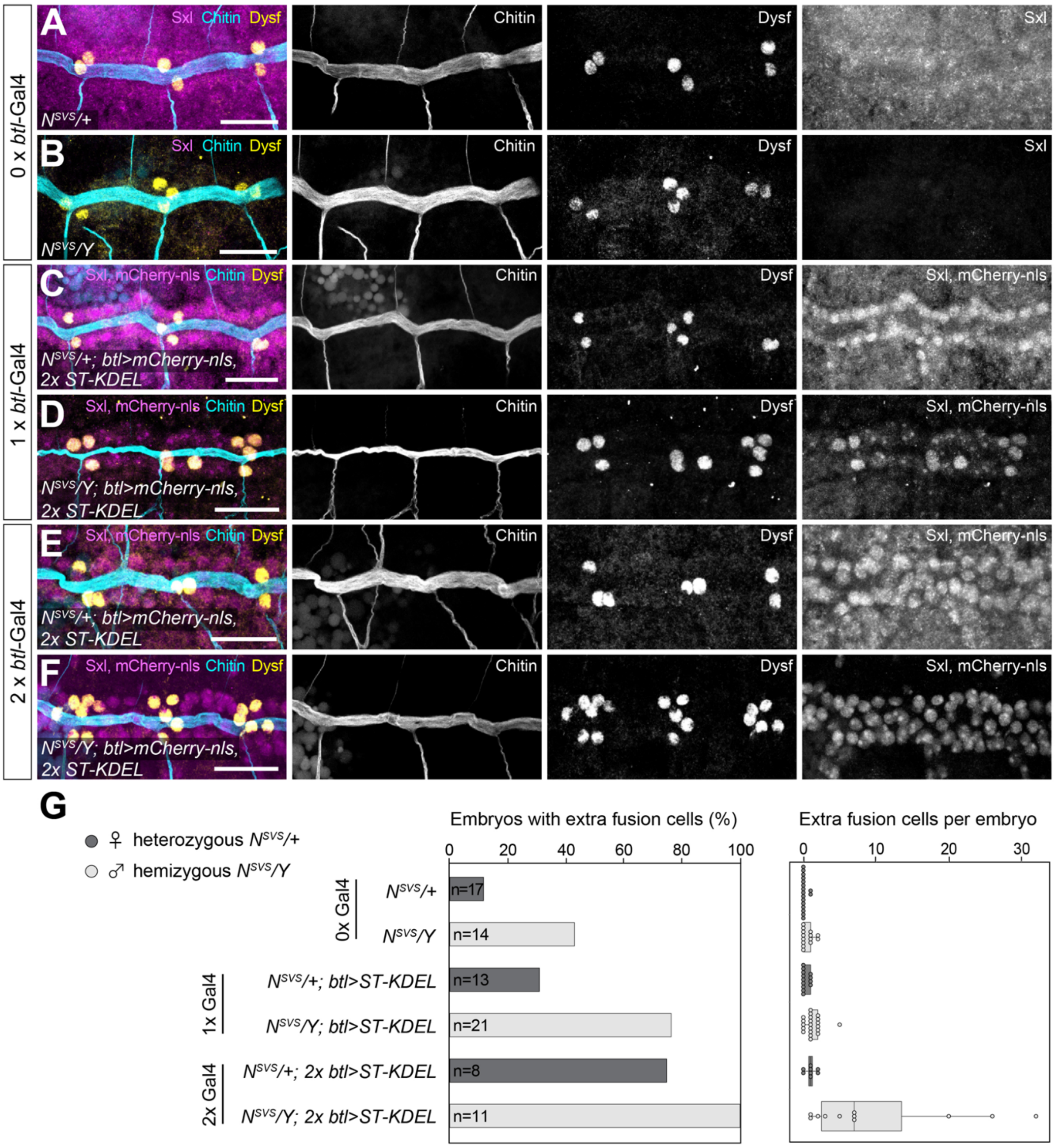


**Figure S6**

**ER retention of Notch::SVS leads to mis-specification of tracheal fusion cells. Related to Figure 7.**

**(A-F)** Representative images of tracheal dorsal trunk in heterozygous *N::SVS/+* (female; A,C,E) and hemizygous *N::SVS/Y* (male; B,D,F) stage 15 control embryos (A,B) or in embryos expressing *ST-KDEL* and *mCherry-nls* under the control of one (C,D) or two (E,F) copies of *btl*-Gal4. Embryos were stained for the female-specific isoform of Sex-lethal (Sxl, magenta), mCherry (magenta), Chitin (cyan) and Dysfusion (Dysf, yellow). Note that elevated levels of ST-KDEL expression driven by two copies of *btl*-Gal4 cause increased prevalence of mis-specified Dysf-positive fusion cells.

**(G)** Quantification of fusion cell specification in the indicated genotypes. The percentage of embryos displaying at least one extra fusion cell (left graph), as well as the number of extra fusion cells per embryo (right graph) increases with *btl*-Gal4-dependent expression levels of ST-KDEL. Sample size (number of embryos, n) is indicated. Boxplot (right graph) shows the median (line), interquartile range (box) and 1.5x interquartile range from the 25th and 75th percentile (whiskers).

Scale bars (A-F): 20 µm.

**Supplemental Movies**

**Movie S1**

**Biotin injection triggers rapid ER release and secretion of Serp-SBG in tracheal dorsal trunk. Related to Figure 3.**

Time-lapse movie of tracheal dorsal trunk in *btl-Gal4 UAS-Serp-SBG, 4xUAS-SA-KDEL* embryo (stage 15) injected with biotin (1 mM) at t=0 min. Gray, Serp-SBG. Time, minutes. Scale bar, 10 µm.

**Movie S2**

**Secretion and re-internalization of Serp-SBG from the tracheal lumen in tracheal dorsal trunk. Related to Figure 3.**

Time-lapse movie of tracheal dorsal trunk in *btl-Gal4 UAS-Serp-SBG, UAS-mRFP-Rab4, 4xUAS-SA-KDEL* embryo (stage 15) injected with biotin (1 mM) at t=0 min. Green, Serp-SBG; magenta, mRFP-Rab4. Arrowheads indicate co-localization of Serp-SBG and mRFP-Rab4. Time, minutes. Scale bar, 10 µm.

**Movie S3**

**Secretion and re-internalization of Serp-SBG from the tracheal lumen in tracheal dorsal branches. Related to Figure 3.**

Time-lapse movie of unicellular tracheal dorsal branches in *btl-Gal4 UAS-Serp-SBG, UAS-mRFP-Rab4, 4xUAS-SA-KDEL* embryo (stage 15) injected with biotin (1 mM) at t=0 min. Green, Serp-SBG; magenta, mRFP-Rab4. Arrowheads at t=25.7 min indicate co-localization of Serp-SBG and mRFP-Rab4. Time, minutes. Scale bar, 10 µm.

**Movie S4**

**The kinetics of Serp-SBG secretion depends on the dosage of injected biotin. Related to Figure 3 and Figure S2.**

Time-lapse movies of tracheal dorsal trunk in *btl-Gal4 UAS-Serp-SBG, 4xUAS-SA-KDEL* embryos (stage 15) injected with biotin of the indicated concentrations at t=0 min. Representative movies are shown. Four injected embryos per biotin concentration were analyzed. Green, Serp-SBG; Time, hours: minutes : seconds. Scale bar, 5 µm.

**Movie S5**

**Transit of Serp-SBG through mCherry-Sec24-labeled ER-exit sites. Related to Figure 4 and Figure S3.**

Time-lapse movie of tracheal dorsal trunk cells in *btl-Gal4 UAS-Serp-SBG, UAS-mCherry-Sec24, 4xUAS-SA-KDEL* embryo (stage 15) injected with biotin (1 mM) at t=0 min. Images were processed using deconvolution. Green, Serp-SBG; magenta, mCherry-Sec24. Time, minutes. Scale bar, 5 µm.

**Movie S6**

**Close-up view of Serp-SBG transit through a single mCherry-Sec24-labeled ER exit site. Related to Figure 4 and Figure S3.**

Time-lapse movie of tracheal dorsal trunk cells in *btl-Gal4 UAS-Serp-SBG, UAS-mCherry-Sec24, 4xUAS-SA-KDEL* embryo (stage 15) injected with biotin (1 mM) at t=0 min. Arrowhead marks a single ER exit site. Images were processed using deconvolution. Green, Serp-SBG; magenta, mCherry-Sec24. Time, minutes. Scale bar, 1 µm.

**Movie S7**

**Dynamics of Gliotactin assembly at tricellular junctions in embryonic epidermis. Related to Figure 6.**

Time-lapse movie of epidermis in embryo (stage 15; *ubi-Dlg1::TagRFP/+ or Y; Gli(CPTI 002805)/2xUAS-ST-KDEL; hh-Gal4 UAS-CyPET-nls/+*) expressing endogenous Gli::SVS (green) and *ubi*-Dlg1::Tag-RFP (magenta). CyPET-nls (cyan) and ST-KDEL (two copies) are expressed in epidermal stripes under the control of *hh-Gal4*. Biotin (1 mM) was injected at t=0 min. After release from the ER, Gli::SVS appears at ERES/Golgi compartments (t=2:00) and subsequently begins to accumulate at tricellular junctions and to a lesser extent at bicellular junctions (t=10:00). Images were processed using deconvolution. Time, minutes: seconds. Scale bar, 10 µm.

**Movie S8**

**Visualization of trafficking of endogenous Fasciclin 2 in follicle epithelium of cultured egg chamber. Related to Figure 5 and Figure S5.**

Time-lapse movie of cultured egg chamber (stage 6) expressing endogenous Fas2::SVS (green) and two copies of ST-KDEL under the control of *GR1*-Gal4 in the follicle epithelium. CellMask-orange (magenta) marks plasma membranes. Biotin was added to the medium at t=0 min (1.5 mM final concentration). Time, minutes: seconds. Scale bar, 5 µm.
